# The Dark Side of Biomolecular Condensates: Quantifying the Role of Nucleic Acids

**DOI:** 10.1101/2025.10.23.683552

**Authors:** Daniele Asnicar, Simone Codispoti, Carlo Morasso, Alberta Ferrarini, Giuliano Zanchetta

**Author notes:** These authors contributed equally (Daniele Asnicar, Simone Codispoti).

## Abstract

Liquid-Liquid Phase Separation (LLPS) is being increasingly recognized as a major organizational principle for proteins and nucleic acids (NAs) in cells, as well as a promising strategy for synthetic biology and biomedical applications. Extensive work has explored the role of protein sequence and properties in the regulation of LLPS. On the contrary, the role of nucleic acids has often been overlooked. Here, to fill this gap we focus on model systems made of oligonucleotides with tuned lengths and degree of hybridization, mixed with a moderately charged, disordered peptide. Combining multiple length scales through experiments and molecular dynamics simulations, we unravel the distinct effect of the properties of NAs on the phase behavior and we propose a metric for the stability of biocondensates. We also characterize the conditions for the onset of liquid crystalline order in the droplets and we show that it is associated with a dramatic slowing down of the dynamics of oligonucleotides, while peptides retain high mobility. Our results can be generalized to natural and non-natural NAs of arbitrary structure, providing a guide for the design of synthetic NA-containing coacervates.

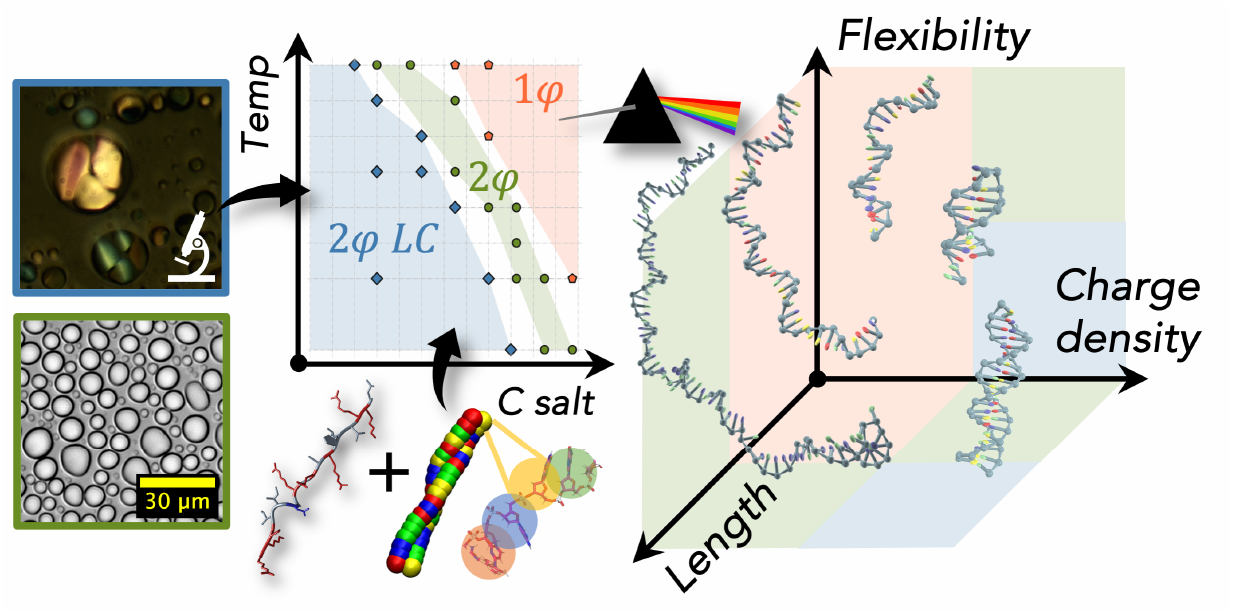

## Introduction

The interactions between proteins and nucleic acids (NAs) are crucial for countless cellular functions, from the assembly and activity of the ribosome [1] to transcription regulation [2]. In particular, NAs can regulate protein solubility and activity through the formation of membraneless organelles, also called biomolecular condensates or coacervates, arising from spontaneous liquid-liquid phase separation (LLPS) of protein-NA complexes [3, 4]. This mechanism, which may have provided enrichment and colocalization in prebiotic times [5, 6], is increasingly recognized as a major principle for cellular organization, from gene expression [2] to DNA repair and response to stress [7, 8], but is also associated with the onset of pathological protein aggregation [9]. Furthermore, LLPS is being investigated as a promising strategy for synthetic biology [10] and drug delivery [11], whose rational design, however, requires robust understanding and fine control [12].

The role of solution conditions (ions, crowders) [13] and of protein sequence and charge patterning in shaping the properties of biomolecular condensates has been widely investigated and is rather well established [3, 4, 9, 14–21]. In contrast, fewer studies have been conducted on the contribution of NAs, often with contradictory observations. Compared to proteins, NAs have less pronounced sequence variability and are characterized by a highly charged backbone, which can overwhelm base composition effects. Overall, purines display higher propensity for phase separation in the presence of positively charged amino acids [22] or divalent ions [23–25], which has been correlated to base stacking free energy [22], direct interaction with amino acids [26] and nucleotide-solvent interactions.

The distinct ability of NAs to base-pairing gives rise to double stranded (ds) constructs which are less flexible, have higher charge density and, by virtue of the lower exposure of the nucleobases, weaker hydrophobic character than single strands. This combination of factors results in a strong effect on coacervation, but it is difficult to unravel each individual contribution [27], also because most studies have been conducted on mixtures of oligonucleotides (ONTs) with highly charged polyK strands [28–31]. The more flexible single stranded (ss) DNA displays coacervate droplets in a wide range of conditions [32]. Instead, dsDNA often forms solid precipitates which dissolve at high enough ionic strength, with isotropic liquid and liquid crystalline droplets at intermediate salt concentrations [29, 30] (in the latter case, the term Liquid-Liquid Crystalline Phase Separation (LLCPS) [33, 34] is used). This general trend agrees with simulation and theory results, showing inhibition of coacervation because of the entropic penalty of aligning rigid poly-electrolyte chains [31, 35] and a weaker degree of structural correlation [36, 37]. However, other studies involving portions of the Histone binding protein report conflicting results, with increased stability for the more highly charged dsDNA and Guanosine (G) quadruplexes [38, 39], or on the contrary enhanced phase separation for (gel-like) ssDNA droplets relative to dsDNA [40]. Even the persistence of the secondary structure inside the condensates is disputed [41, 42].

Here, we systematically explore the phase behavior of NA oligomers of different lengths and secondary structures, including partially structured systems (Figure 1A), when mixed with a disordered peptide carrying a moderate positive charge and without pronounced sequence patterns (Figure 1B). This condition favors a more realistic comparison with naturally-occurring proteins, better mimics the NA interactions within biomolecular condensates [28] and allows the differences between NA structures, often leveled out by peptides with a high charge density, to emerge more clearly.

**Figure 1:**
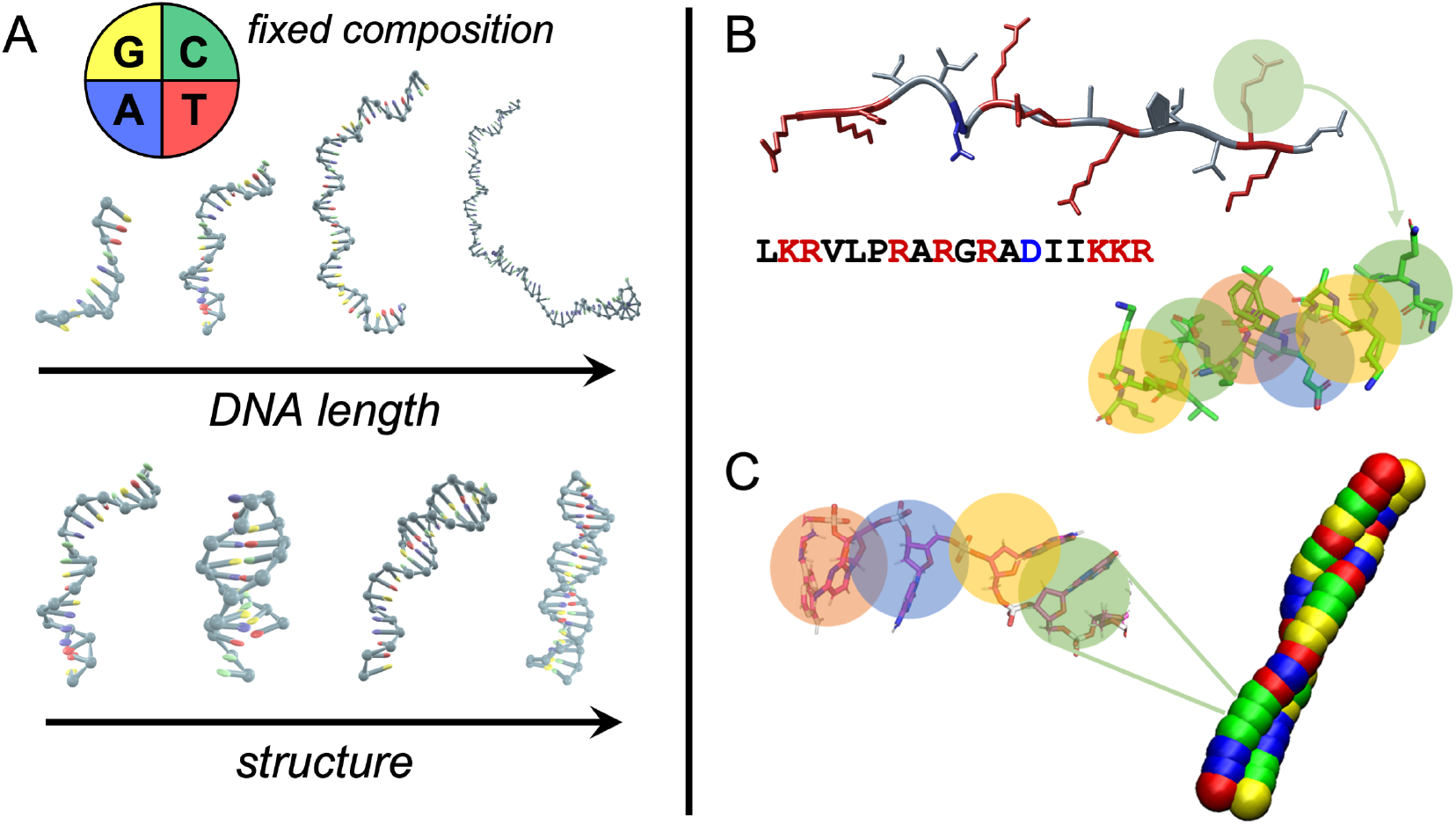
Overview of the systems investigated in this study. A) oxDNA sketches of DNA oligomers of balanced NTs composition, including single strands of different lengths, along with fully and partially base-paired strands (see Table S2). B) Structure and sequence of the pL22 peptide (neutral amino acids are indicated in black, cationic AAs in red, anionic AAs in blue) and corresponding CG model, where each bead represents an amino acid. C) CG model of dsDNA adopted in MD simulations, with one-bead representation of nucleotides.

The experimental investigation is critically supported by simulations based on a coarse grain (CG) model with resolution at the level of a single nucleotide/amino acid and parameters tuned to capture sequence-specific interactions and mechanical properties of the polyelectrolytes [16, 43, 44], which we adapt to include hybridization. This kind of models cannot provide an explicit description of phenomena such as solvent reorganization, salt partitioning or counterion release upon binding [45–47]. Nevertheless, they have been found to be able to capture the main effects of the phase behavior of protein-NA mixtures, so greatly contributing to our understanding of LLPS and material properties of biomolecular condensates [39, 48, 49].

We establish robust quantifiers of flexibility and charge distribution for the various ONTs of different lengths and secondary structures and dissect their role in coacervate stability. CG modeling also allows us to map the interaction network and mobility of biomolecular species inside the coacervates, providing molecular level understanding about the onset of liquid crystalline ordering.

We extend the investigation to Peptide Nucleic Acids (PNA) and polyphosphate (polyP), which, besides being instrumental to our analysis, have on their own biological and biotechnological interest. In particular PNAs, with nucleobases attached to a neutral, peptidic backbone, also thanks to their enhanced stability and specific pairing to DNA and RNA [50], are a powerful tool for diagnostics, gene editing and drug delivery [51]. Polyps, which instead have similar backbone to DNA and RNA but lack nucleobases, are ancient and ubiquitous inorganic polymers, involved in the structure and function of chromatin [52], with important biomedical applications [53]. Their propensity to form condensates has recently attracted attention in the context of prebiotic research [54] and cell biology [55, 56], where polyP may modulate gene expression [57, 58]. Within our framework, we model and explain the phase behavior of such biomolecular species.

## Results and discussion

To decouple the role of the molecular features of NA from the sequence composition, we select DNA strands with fixed 25% fraction for every nucleobase. We explore DNA lengths from 10 to 80 NTs and we design and permutate sequences to tune the degree of intra- and inter-strand hybridization, from ss to hairpins to full ds (see Figure 1A). The complete set of DNA oligomers employed in this study is reported in Table S2. We evaluate their behavior in solution with the pL22 peptide from *Thermus thermophilus*’ ribosome (see Figure 1B and Table 1). The choice of this peptide is motivated by its average composition, with polar, apolar and hydrophobic amino acids, and a moderate fraction of charged residues. Indeed, its low *Sequence Charge Decoration* (SCD) value [59], resulting from interspersed positively charged LYS and ARG residues and from the presence of a negatively charged ASP, makes it a more realistic representative of DNA-binding proteins (see Fig. S1) than the typically used polyK, and thus better suited to study the effects of DNA structure on coacervation. Moreover, pL22 being a protein fragment buried in the large ribosomal subunit, it naturally interacts with RNA and its sequence is highly conserved across the *bacteria* and *archaea* domains [5].

**Table 1:** pL22 sequence and principal physicochemical properties. Positively charged residues at neutral pH are highlighted in red, negative ones in blue; “GRAVY” stands for “Grand Average of Hydropathy, “SCD” for Sequence Charge Decoration.

|  |  |
| --- | --- |
| Sequence | LKRVLPRARGRADIIKKR |
| Molecular weight | 2146.6 Da |
| Net charge at pH 6-8 | + 7.0 |
| Theoretical pI | pH 12.58 |
| SCD | + 2.8 |
| GRAVY | −0.85 |

### Coarse grain modeling of biomolecular interactions

To map and analyze the experimental phase behavior, we perform molecular dynamics (MD) simulations based on a CG model previously developed for proteins [16, 43] and for RNA [44]. Each amino acid or nucleotide is represented by a bead (see Figure 1C), with interaction parameters that reflect the hydrophobicity and ionic charge of the various moieties, whereas the solvent is implicit and small ions are taken into account through the Debye-Hückel approximation (more details in SI). Here, we introduce into the model hybridization (see Materials and Methods), which is crucial to catch the effect of the secondary structure of ONTs. Thus, the chemical and structural specificity of the model allows us to directly compare the numerical predictions with the experimental results.

### Characterization of the phase behavior

The stability and properties of NA/protein condensates are known to be affected by the polyelectrolyte charge ratio [39, 48, 60]. To properly compare the behavior of NAs of different lengths and structures, we therefore consider electro-neutral mixtures. While both DNA oligomers and peptides are fully soluble in the buffer as single species, their mixtures spontaneously separate, already at ∼500 *µ*M overall concentration, into concentrated, liquid-like droplets, enriched in both DNA and peptides, in coexistence with a dilute supernatant (see Figure 2A). The onset of condensation is also observed in the CG simulations, as reported in Figure 2B. The phase boundaries and the concentration of DNA and peptide within the phases depend on the total concentration, on the temperature *T* and on the total ionic strength ℐ 𝒮 of the mixtures. A typical bell-shaped phase boundary in the concentration-ionic strength plane is observed [3] (Fig. S2). For a given concentration, both increasing temperature and ionic strength destabilize the condensed phase, as shown in Figure 2C for a ssDNA 10mer, for experiments (*top*) and simulations (*bottom*). The two plots show remarkable agreement: the biphasic region appears only for ℐ 𝒮 values below 100 mM and narrows with increasing temperature. Compared with previous studies of DNA oligomers with highly charged polycations, such as polyK and polyR, in which LLPS up to ℐ 𝒮∼800 −900 mM was reported[28, 29], the system exhibits a salt resistance much closer to physiological levels, a behavior that can be attributed primarily to the more distributed charged units in pL22, similar to natural proteins.

**Figure 2:**
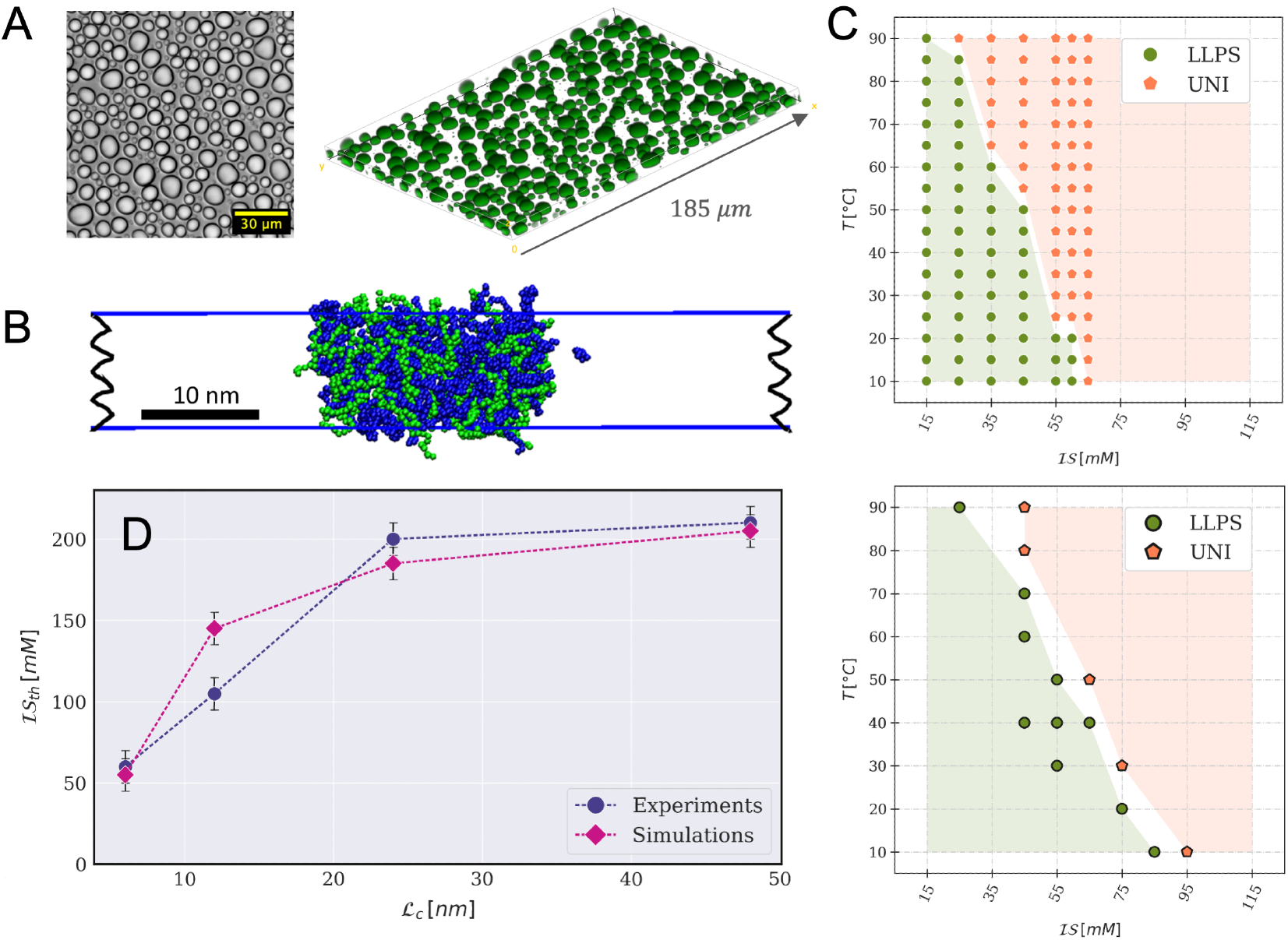
Characterization of the phase behavior of peptide/ssDNA mixtures. A) Widefield microscopy image of pL22/ssDNA-20 coacervates (at ℐ 𝒮 =15 mM and T =298 K) sedimented on the substrate (left); confocal 3D-slab reconstruction (right) of the same sample with 1:200 FITC-labeled DNA strands. B) Snapshot from a CG-MD slab simulation of pL22/ssDNA-20 (blue and green, respectively) at ℐ 𝒮=75 mM and T =303 K. C) Experimental (top) and simulated (bottom) phase diagrams of pL22/ssDNA-10, as a function of the ionic strength ℐ 𝒮 and of the temperature T ; stoichiometric concentrations are: 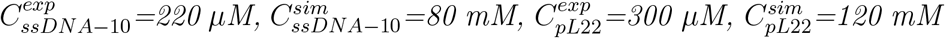. Green full circles and orange pentagons denote bi- and mono-phasic states, respectively, observed in the sampled conditions. Green and orange shades indicate the extent of the two regions. D) Highest ionic strength at the boundary of the LLPS region for T =298 K observed in experiments (blue) and simulations (violet), as a function of the contour length ℒ_c_ of ssDNA strands with 10, 20, 40 and 80 NTs.

To characterize the coacervation propensity for each NA, we can compare their highest ionic strength at which the biphasic region is observed at room temperature, ℐ 𝒮_*th*_. Although this value does not correspond to the true critical point located at the maximum of the spinodal line, it is a good proxy for it, because the topology of the phase boundaries remains unaffected [22].

### Effect of DNA length and secondary structure

The analysis of systems that differ only in the length of ssDNA allows us to dissect the contribution of NA chain connectivity in condensate formation separately from the electrostatic interaction between NA-peptide monomeric units. In Figure 2D, we plot ℐ 𝒮_*th*_ as a function of the contour length ℒ_*c*_ of ssDNA filaments (the experimental and simulated phase diagrams of ssDNA-20 are shown in Figure 3, while the other related diagrams are reported in SI, Fig. S3). In both experiments and simulations, in agreement with previous reports on different NA/peptide systems [39, 40, 48, 61, 62], we first observe an increase of ℐ 𝒮_*th*_ for short ONTs, which then levels off beyond ssDNA-40. We verify that a scrambled version of ssDNA-20, retaining unpaired conformation, has very similar behavior (data not shown). For both neutral polymers and mixtures of oppositely charged polyelectrolytes, an increase in chain length favors phase separation due to the decrease of translational entropy penalty [36, 63, 64]. Such growth and saturation can be qualitatively described within the framework of the mean-field theory of Overbeek and Voorn [65].

**Figure 3:**
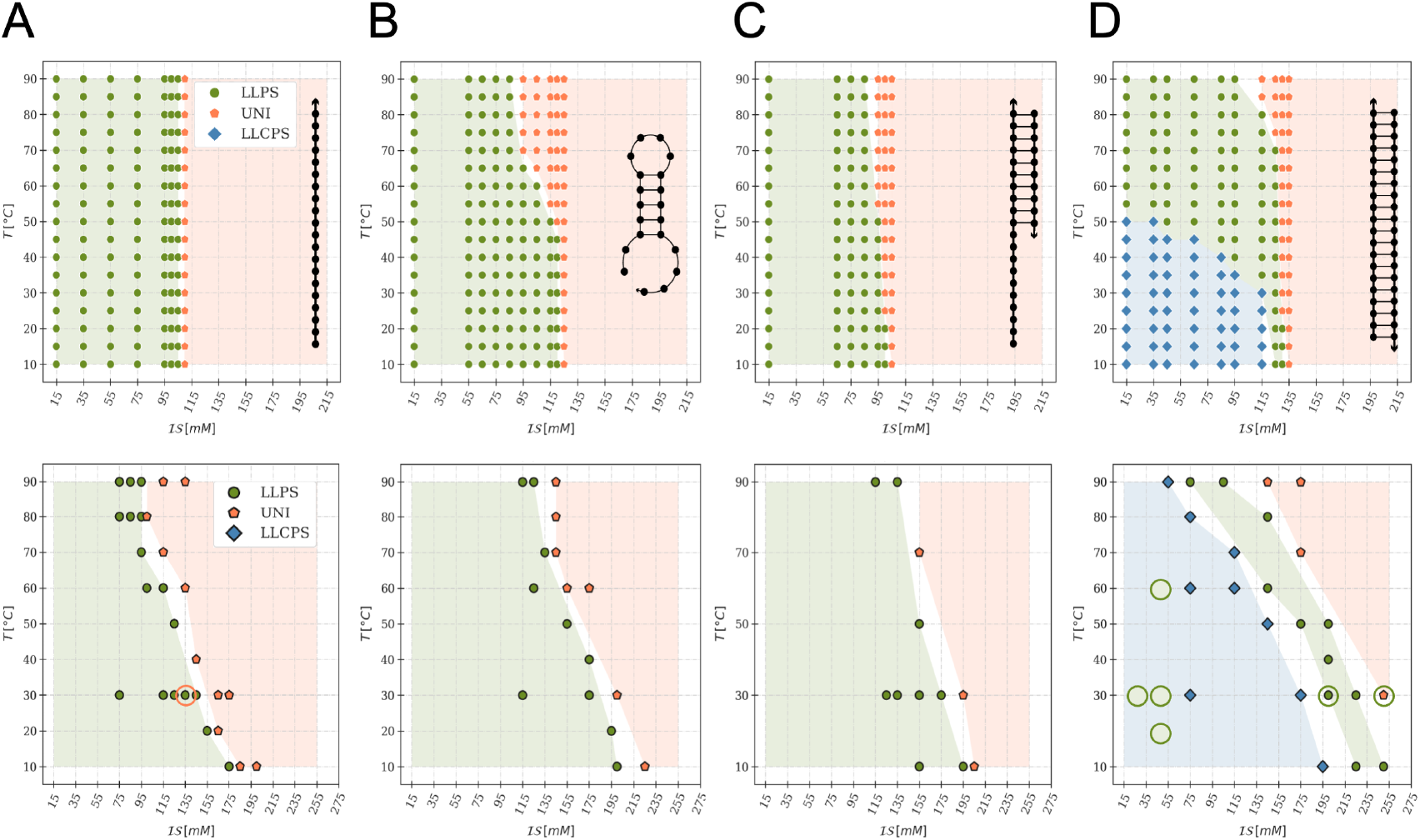
Experimental (top) and simulated (bottom) phase diagrams of mixtures of pL22 with oligonucleotides with different degrees of hybridization, as sketched in each panel. (A) 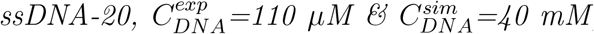, (B) 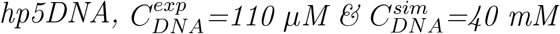, (C) 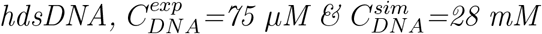 (D) 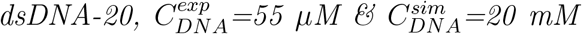. In all cases, 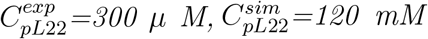, so to achieve overall charge neutrality of the mixtures. The phase behavior is denoted by the color and shape of markers and shaded regions: monophasic (orange pentagons), isotropic biphasic (green full circles) and liquid-crystalline biphasic (blue diamonds). The orange (green) empty circles in the simulation diagrams of A and D denote states sampled using a higher (lower) bending constant values in the oligomer model (Eq. S6, see text for details).

Next, to investigate the role of DNA secondary structures in LLPS, we progressively introduce ds portions into the filaments, without affecting the base composition. Figure 3 shows the measured and simulated phase diagrams for mixtures of pL22 with ssDNA-20 (A); with hp5DNA, a 5 bp-stem hairpin obtained by reshuffling the sequence of ssDNA-20 (B); with hdsDNA, a half-structured ONT, i.e. a ds 10mer with an unpaired tail of 10 NTs (C); and with fully base-paired dsDNA-20 (D). Compared to single filaments, base-paired structures are stiffer, with a much higher persistence length (∼50 nm for dsDNA vs ∼0.75 nm for ssDNA at 10 mM Na^+^) [66]. They also have fewer exposed aromatic nucleobases, which can engage in short-range interactions with the peptide. Both features would be expected to destabilize the coacervates for ds structures [35–37]. On the other hand, previous experiments on polyK mixed with partially or fully hybridized DNA structures (with significant variations in base composition) [28, 29] found a sharp transition from liquid coacervates to solid precipitates whenever the ds portion exceeded 40%. This behavior was ascribed to the higher charge density of ds structures, interacting with the highly charged polyK. Partially hybrid structures, such as hp5DNA, could combine the advantages of both high charge density and relatively high flexibility, as was suggested by the observation of particularly high resistance to their release from complexes with polyK [67].

With the weakly charged pL22 we observe LLPS in all of our samples, with an overall increase in stability from ssDNA-20 to dsDNA-20. This confirms the important role of charge density, which, however, was never quantitatively addressed. We will discuss this point in the next Section.

In addition to its higher ℐ 𝒮_*th*_ for LLPS, dsDNA-20 differs from less structured DNA strands in the appearance of a liquid crystalline phase within the demixed droplets (light blue region in Figure 3D) [29, 30, 33]. This ordered yet fluid phase transitions to the isotropic liquid state at temperatures above ∼50^°^C or close to the ionic strength threshold. ONTs and peptides are colocalized in droplets, as assessed by Raman microscopy (Figure 4A,C): the intensities of DNA- and peptide-specific bands in the Raman spectrum [68, 69] (Figure 4G) display a correlated, non-uniform spatial distribution of the two biomolecules. Moreover, the anisotropic arrangement of DNA helices, a hallmark of LC phase, is evidenced by droplet birefringence (Fig. S4) and by the anisotropic fluorescent emission of an intercalated dye, SYBR green. In Figure 4B, the confocal microscopy image displays alternating bright and dark regions within the fan-shaped domains (suggesting a columnar phase) because the horizontally polarized incident laser has maximum (minimum) efficiency of excitation for horizontal (vertical) dye molecules, corresponding to vertically (horizontally) lying dsDNA helices. In contrast, no fluorescence anisotropy is observed for dye-labeled peptides (Figure 4D), a sign of their overall disordered arrangement.

**Figure 4:**
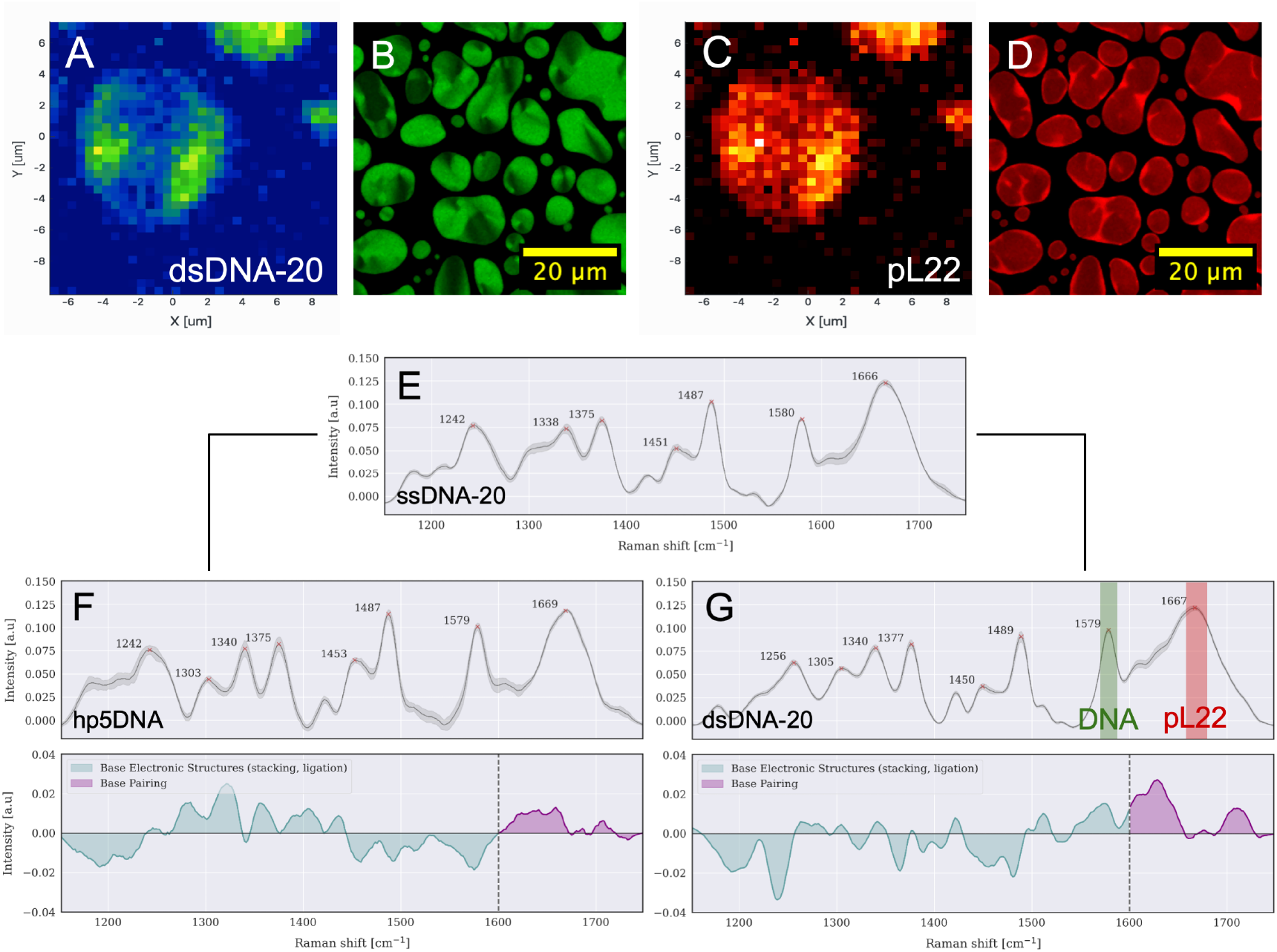
Experimental characterization of molecular arrangement and secondary structure inside the condensates. A-C) Ramam microscopy images of a coacervate droplet of pL22/dsDNA-20 (C_DNA_ =55 µM, C_pL22_=300 µM, ℐ 𝒮=15 mM, T =298 K). The color scales correspond to the integrated intensity of the two bands highlighted in the spectrum of dsDNA (panel G), around 1580 cm^−1^ for DNA ring stretching (green) and around 1660 cm^−1^ for peptide Amide I (red). B,D) Confocal images of pL22/dsDNA-20 coacervates (C_DNA_=55 µM, C_pL22_=300 µM, ℐ 𝒮=75 mM, T =298 K). SYBR Green intercalating dye marks the anisotropic arrangement of DNA strands (B), while a fraction of peptides are labeled with TAMRA (D). E) Average Raman spectra and variations inside the droplets of pL22 (C_pL22_=300 µM) with ssDNA-20 (C_DNA_=110 µM). F,G) Average Raman spectra and variations inside the droplets of pL22 (C_pL22_=300 µM) with hp5DNA (C_DNA_=110 µM) (F) and dsDNA-20 (C_DNA_=55 µM) (G) atℐ 𝒮 =154 mM and T =298 K. The differences between pairs of spectra (last row) indicate in both cases that DNA strands in the coacervates are hybridized.

The existence of LC order clearly reflects the stiffness and anisotropy of dsDNA: it is well known that in bulk, under suitable conditions, long [70] and shorter [71, 72] dsDNA, like other sufficiently stiff biopolymers [33], can organize into ordered fluid phases, driven by excluded volume effects of DNA strands and their aggregates [71, 73], or by counterion-induced attraction [72]. Here, the cationic peptide promotes both demixing and ordered self-assembly [30, 33].

The experimentally observed trends of the mixtures of pL22 with ONTs of different secondary structures are confirmed by the analysis of the simulated phase diagrams (bottom row in Figure 3). In particular, dsDNA-20 exhibits the same sequence of LC and isotropic liquid regions at increasing ℐ 𝒮and *T*. The calculated spatial density distribution and translational diffusion coefficient demonstrate the (isotropic or anisotropic) liquid nature of the concentrate phase, distinctly different from those computed in the presence of polyK (Fig. S5). Despite their overall similarity, one can notice that the stability of the LLPS region is overestimated in the simulated phase diagrams as compared to the experimental ones, in particular for the LC region of dsDNA. One possible source for the discrepancy is that the total polymer concentration in the simulations is significantly higher than in the experiments, for computational reasons; therefore, the boundary of the two-phase region is crossed at a different ionic strength (Fig. S2) [22]. In fact, simulations performed at the experimental concentration showed a decrease in ℐ 𝒮_th_ (Fig. S6); however, very small aggregates form under such conditions, which makes the systematic investigation of phase separation more uncertain, and therefore we opted for higher concentrations.

The systematic over-stability of simulated phase diagrams may also derive from the limitations of our CG model. In particular, the same parameterization for NT-AA interactions is used for paired and non-paired nucleotides, that is, for dsDNA and ssDNA. As a consequence, reduced exposure of the bases in structured DNA is ignored [74], which may lead to an overestimate of hydrophobic interactions, particularly relevant for dsDNA and at higher ionic strengths.

Moreover, some discrepancy between the experimental and simulated phase boundaries may be attributed to partial unfolding of hybridized ONTs in the experimental samples [42]. Indeed, while hp5DNA is stable up to around 50 ^°^C, hdsDNA melts around 40 ^°^C (Fig. S7). However, Raman spectra highlight the presence of base pairing and stacking signatures within the coacervates at room temperature [68, 69] (Figs. 4 and S8): the average intensities in the 1150-1750 cm^*−*1^ interval show significant differences for both hairpins and double helices with respect to the corresponding signals from ssDNA (last row in panels F and G, respectively), confirming that the ONTs mostly retain their hybridized conformations.

### Charge heterogeneity and flexibility determine the stability of condensates

In order to describe the observed trends in the phase behavior of various ONTs and connect them to their single-chain properties, we must properly address the molecular features of ONTs: while length is a well-known determinant of phase behavior of DNA and polyelectrolytes in general [62], flexibility and charge density have opposite effects, but the quantities typically used to measure them can defy intuition, are often interconnected, or fail to capture subtle yet important differences. We briefly introduce here our approach to define simple but meaningful parameters to describe the critical NA properties that determine their coacervation.

#### Length

Because we compare the behavior of ss and ds ONTs, or their combination, the number of nucleotides is not an ideal metric. To consider the different building blocks, we rather consider the contour length ℒ_*c*_ of the various DNA strands, defined as the sum of ss and ds steps, of length 0.6 nm and 0.34 nm, respectively [75].

#### Flexibility

Within the CG models, flexibility is conveniently tuned through a bending potential between adjacent beads in a strand, independent of temperature and ionic strength (Eq. S6), and this contribution can have a significant effect on the phase boundaries [35, 37]. For ssDNA, bending is typically neglected, but adding a bending potential reduces the width of the LLPS region (orange empty circle in Figure 3A-*bottom*, obtained with the bending term *k*_b_ = 2 kcal/mol rad^2^).

Likewise, stiffness is easily defined in long, semi-flexible polymers in terms of persistence length, i.e. the typical correlation length of the direction of polymer axis, but in oligonucleotides the very definition of persistence length becomes meaningless, especially for partially folded structures. The related concept of Kuhn length, which represents the length of each segment of a freely-jointed chain, also fails to describe short, heterogeneous strands and results depend on the ONT length [76]. However, the definition of a parameter to characterize the flexibility of real polymeric systems with different architectures is far from obvious. A simple bending rigidity parameter cannot take into account secondary structure and sample conformational fluctuations in an effective way. We find that a suitable quantity able to capture the overall flexibility is the Root Mean Square Deviation (RMSD) of the atomic positions from their average value, normalized by the number of monomers to decouple the residual linear length dependence (Eq. 2 and Fig. S10). In practice, for NAs we use the coarse grained representation provided by *oxDNA* (see Materials and Methods) (Fig. S9).

#### Charge distribution

It is quite natural to think that dsDNA has a higher charge density than ssDNA because the two strands are paired, which in turn strongly affects their arrangement and stiffness. However, a quantitative and applicable definition of charge density requires determining an appropriate length, surface, or occupied volume, which is challenging for heterogeneous structures with spatially distributed charges. We find that the average electrostatic potential |Φ_*e*_|, experienced by a probe charge moving along the surface of an atomistic representation of the ONT, being sensitive to the spatial distribution of charges, can capture such effects. To take into account the intrinsic dependence of the average electrostatic potential on the polymer length (see SI), we use the normalized quantity 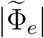 (Eq. 4).

In Figure 5 we plot the experimental ℐ 𝒮_*th*_ as a function of the 3 selected parameters (panel A) and with the separate dependence on (B) *RMSD*_*n*_, (C) 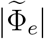 and (D) ℒ_*c*_. In the latter panels, the various colors correspond to different structures. ss and ds families (red dots and orange diamonds, respectively) display clearly different values of flexibility and electrostatic potential, almost independently of their lengths, while partially hybridized ONTs have intermediate values.

**Figure 5:**
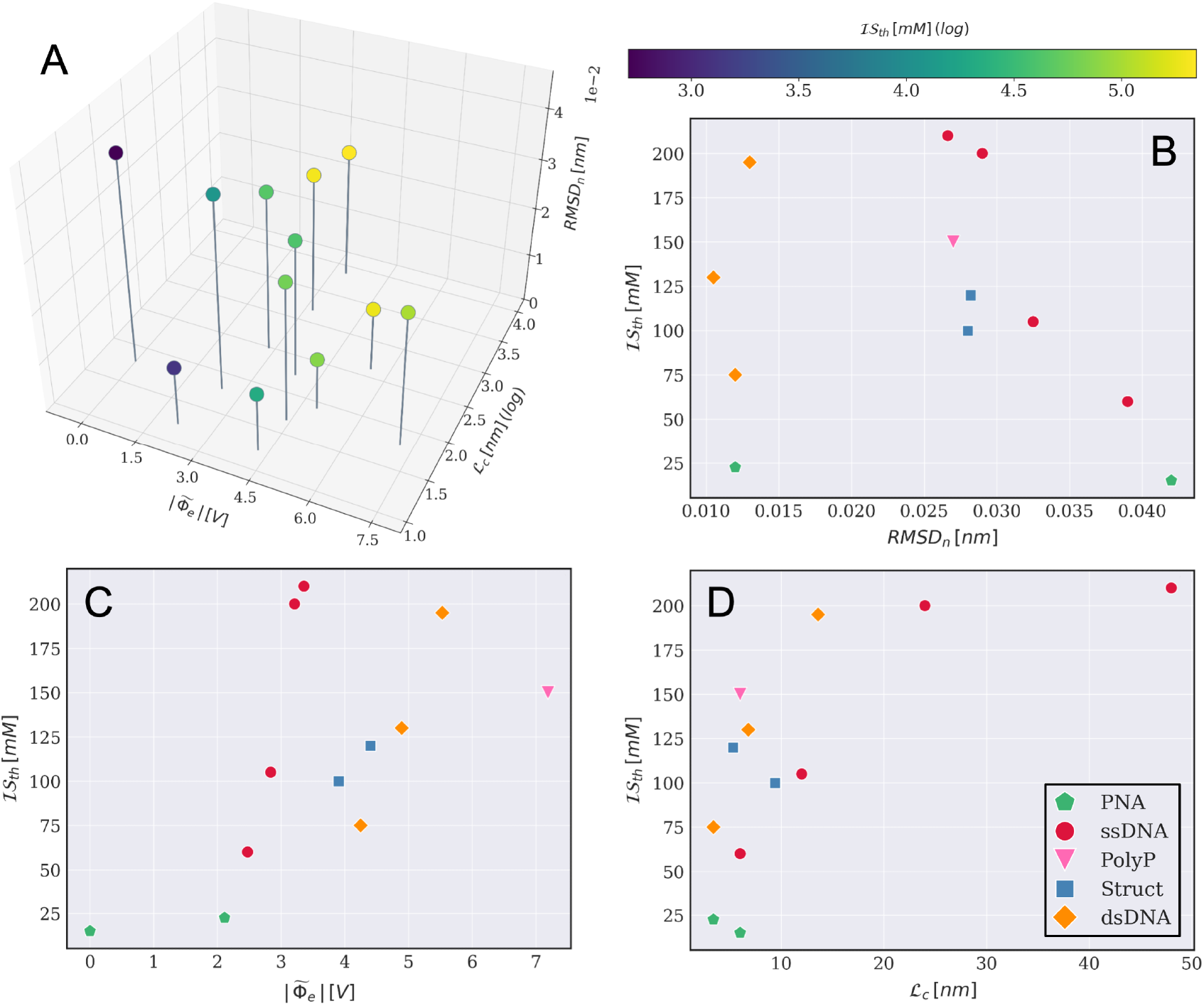
Experimental ionic strength threshold ℐ 𝒮_th_ at the LLPS boundary at 300 K, for mixtures of pL22 with ONTs, PNA and polyP, as a function of the contour length ℒ _c_, of the normalized average electrostatic potential 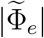 and of the normalized RMSD_n_ of the polyanions. (A) 3D plot with color-coding of the ionic strength (color bar in log scale, top-right). (B-D) Dependence of ℐ 𝒮 _th_ on: (B) RMSD_n_, (C) 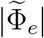 and (D) ℒ _c_. Colors and markers corresponds to different families of structures, as indicated in the caption.

The strongest dependence of ℐ 𝒮_*th*_ is found for length (Figure 5D): similarly to the ss family already discussed above, the LLPS stability for ds ONTs increases with increasing contour length, but with systematically higher ℐ 𝒮_*th*_ values than the more flexible - but less charged - ss ONTs of similar contour lengths.

Indeed, despite previous reports on ONTs with varying sequence composition [27, 36, 37], our experiments suggest that, for DNA oligomers in the 10-100 nt range, the flexibility of the strands alone does not play a major role in determining the stability of the coacervates. On the contrary, larger 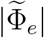 values positively correlate with enhanced LLPS (Figure 5C). The subtle trade-off between the two parameters can be better appreciated in the partially hybridized structures hp5DNA and hdsDNA, which show a similar salt resistance to ssDNA-20 and intermediate between dsDNA-10 and dsDNA-20. In particular, the hairpin structure, besides being more stable than independent filaments, combines highly flexible tracts with large charge density and thus maximizes LLPS propensity. It should be noted that, although in simulations it is possible to tune flexibility without affecting charge density (Fig. 3A), in real DNA systems some degree of coupling between the two parameters always takes place.

### Beyond nucleic acids

The overall trend summarized in Figure 5A suggests that LLPS is enhanced for longer strands and, for a given contour length, electrostatic interactions prevail over flexibility. To further test the generality of this description, we extend our investigation to NA variants, separating the contribution of the charged backbone from the base pairing moieties and better decoupling flexibility from charge density. To this end, we mix pL22 with (i) polyphosphates, as a model for the NA backbone [52, 54], (ii) PNA filaments, which combine an uncharged peptidic backbone with nitrogen bases, and (iii) hybrid ds helices formed by a DNA strand and a complementary PNA filament [50], with flexibility similar to dsDNA but half its charge content (Figure 6C).

**Figure 6:**
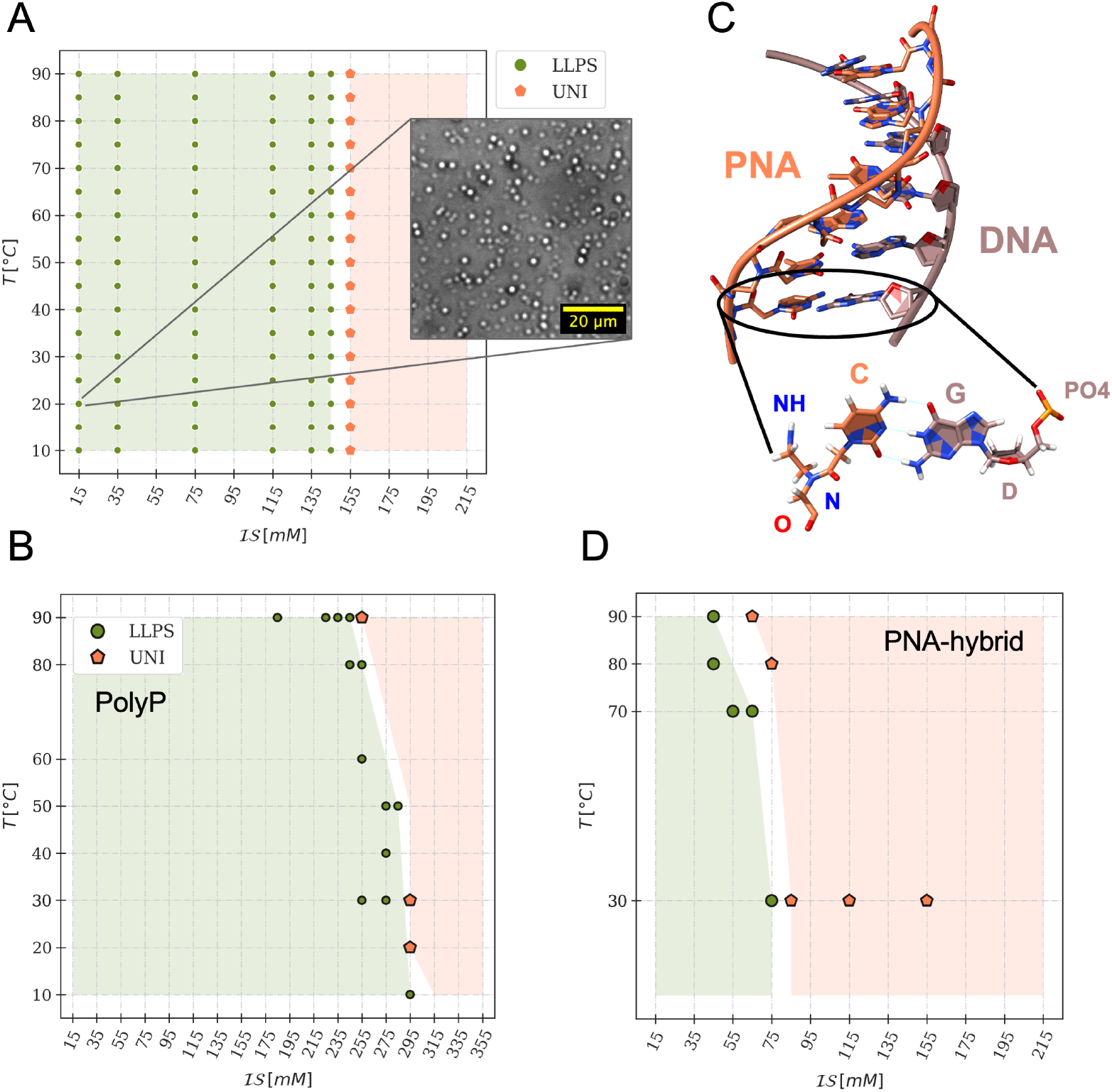
Coacervation of non-standard NAs strands. A,B) Experimental (A) and simulated (B) phase diagrams of pL22/polyP mixtures at stoichiometric concentrations 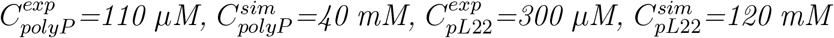. The phase behavior is denoted by the color and shape of markers and shaded regions: monophasic (orange pentagons), isotropic biphasic (green full circles) and liquid-crystalline biphasic (blue diamonds). The inset in A shows a widefield micrograph of the system at low ionic strength and ambient temperature. C) Structure of a PNA-10 hybrid (PDB 1PDT); as the C-G WC pairing shows, PNA and DNA share canonical nucleobases, but the deoxyribose-phosphate backbone of DNA is replaced by a pseudopeptide skeleton in PNA. D) Simulated phase diagram for mixtures of pL22 (C_pL22_=120 mM) with the PNA-20 hybrid (C_PNA−20_=40 mM).

While the neutral PNA does not display LLPS at any condition, a polyP 20mer forms liquid droplets following the general trend found for DNA oligomers (Figure 5A), but with strikingly different properties. Indeed, polyP features both high charge density and high flexibility and was previously shown to phase separate in mixtures with other positively charged peptides and proteins [54–56, 77]. Interestingly, we find that it displays higher salt resistance than dsDNA-20, but with markedly smaller droplets both at low and high ionic strength (Figure 6A). Simulations based on a CG model of polyP, built in analogy with the model used for DNA (see Materials and Methods), also predict high stability and confirm the major role played by the charge density (Figure 6B). However, the threshold ionic strength is largely overestimated as compared to experiments, which points to the need of a more specific parameterization for quantitative agreement.

As regards DNA-PNA hybrids, experimental investigation is only possible for 10mers, because of solubility limits of longer strands: no LLPS is observed at concentrations and solvent conditions similar to the ones tested for ssDNA-10 and dsDNA-10 (see SI, Fig. S12). However, simulations are extended to a 20mer to directly compare it to its ss and ds DNA counterparts (Figure 6D). For the PNA strand, the same CG model of ssDNA is used, with charges set equal to zero. The DNA-PNA hybrid exhibits lower ℐ 𝒮_*th*_ than ssDNA, which has a very similar charge density but is much more flexible. As expected for its reduced electrostatic potential, it is also much less stable than dsDNA. Moreover, LC ordering is suppressed for pL22/PNA-20 hybrid.

Based on the experimental and simulated trends for DNA and single NA components (Figures 5A and 6), it would be possible to assess the boundaries of the expected LLPS behavior for other NAs canonical and non-canonical forms, for which we can estimate ℒ_*c*_, *RMSD*_*n*_ and 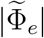 (some of such systems are reported in SI, Fig. S11). For example, for a given ℒ_*c*_, the *RMSD*_*n*_ and 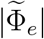 of an RNA single strand are comparable to ssDNA, which is consistent with similar reported phase behavior in the literature [25, 48, 78]. Instead, other folded structures such as G-quadruplexes, which are much stiffer and have a larger electrostatic potential than dsDNA, have been shown to display higher condensate stability, while easily giving rise to solid aggregates [38].

### The onset of liquid crystalline ordering

dsDNA endows coacervates with a distinct sensitivity to the state variables, which is tuned by the polycations. In mixtures of dsDNA oligomers with highly charged amino-based polycations or peptides, liquid coacervates at high NaCl concentration (≳ 500 mM) transform into precipitates at low ionic strength [28]. In mixtures with polyK [29, 30] a richer phase diagram was reported, including various LC phases, again with the isotropic liquid phase only present at high salt concentration (≳ 800 mM) or high temperature. Here, with pL22 we detect LC ordering in a wide range of temperatures and nearly physiological ionic strengths (see Figures 3D and 4B,D); unlike the previous studies, at room temperature the isotropic phase appears already at a relatively low ionic strength, ℐ 𝒮 ∼100 mM.

Previous generic models of charged polyelectrolytes [79–81] found the occurrence of a first order isotropic-to-LC transition inside coacervate droplets, driven by short-range anisotropic (excluded volume) interactions and by electrostatic interactions, if at least one of the polyelectrolytes has weak flexibility and a sufficiently high aspect ratio. Previous slab simulations of dsDNA-containing coacervates [39] missed the onset of LC ordering, probably because of a different modeling of oligonucleotides; in fact, stiffness and aspect ratio of the latter turn out to be crucial to this purpose.

Remarkably, we find the LC phase in our CG simulations (Figure 3D). Figures 7A,B show two configurations, extracted from MD trajectories, of peptides and NAs in isotropic condensates formed by ssDNA-20 and in LC condensates formed by dsDNA-20. While rod-like ds are on average aligned along a common direction within the LC phase, coiled ss are randomly oriented in space. Surprisingly, such disparity does not appear for the peptides, where we cannot detect a significant degree of order in either case.

**Figure 7:**
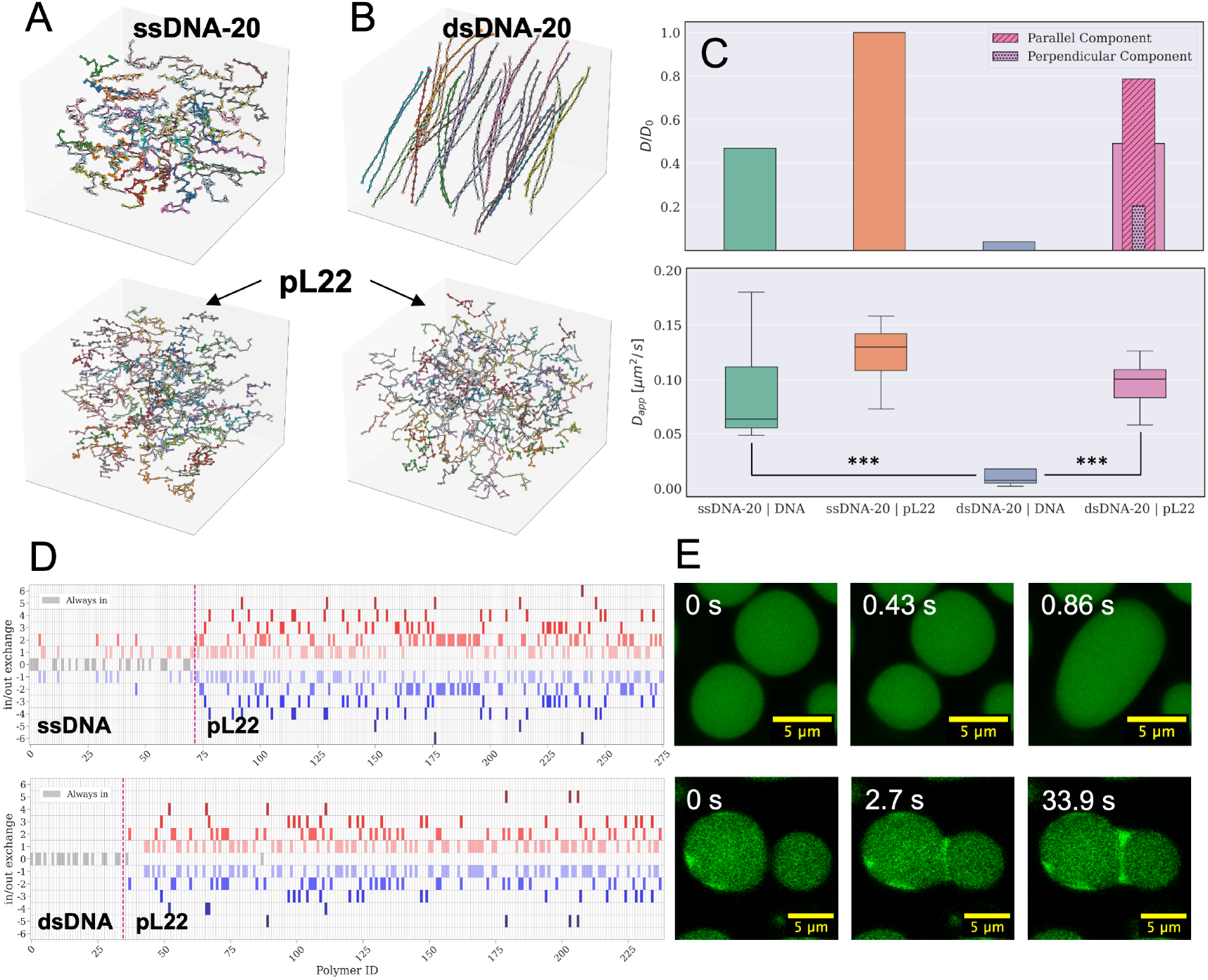
Ordering and dynamics of DNA and peptides inside isotropic and liquid crystalline condensates. A,B) MD simulation snapshots of molecular distributions for DNA oligomers (top) and peptides (bottom) in mixtures of pL22 with ssDNA-20 (A) and dsDNA-20 (B). C) Diffusion coefficients of pL22 and DNA oligomers extracted simulations (top) and from FRAP measurements (bottom) on isotropic condensates of ssDNA and LC condensates of dsDNA; simulation data in the barplot are rescaled by D_0_, the diffusion coefficient of pL22 in isotropic condensates. For pL22 in LC condensates, the contributions from the perpendicular and parallel directions of motion with respect to the director are shown and marked with different hatches and colors. The experimental boxplot results from the fitting of 10 ≤ n ≤ 14 independent recovery curves from different droplets for each of the two samples (ssDNA-20 & dsDNA-20) and two channels (pL22-TAMRA & DNA-FAM). The *** denotes a p-value less than 0.001 in a Mann-Whitney-U two-sided test. D) Number of in (positive)/out (negative) exchanges from clusters, calculated for DNA oligomers (left part) and peptides (right part), during the last 3 µs of MD trajectories of pL22/ssDNA-20 (top) and pL22/dsDNA-20 (bottom). E) Confocal images of fusion events for droplets of pL22 and fluorescently labeled ssDNA-20 (top) and dsDNA-20 (bottom1)8. All the MD results are for simulations at ℐ 𝒮 =75 mM and T =303 K, matching the experimental conditions of ℐ 𝒮 =75 mM and T =298 K.

Further quantitative insight on the molecular organization inside the coacervates is provided by the radial distribution function (rdf) for the amino acids and the NTs in different mixtures, quantifying the degree of spatial correlation of the different species (see SI, Fig. S5). In isotropic coacervates (ssDNA and dsDNA at high) rdfs indicate direct contacts between amino acids and nucleotides, but no contacts between the nucleotides and a few contacts between peptides. Instead, in LC coacervates of dsDNA at low, the rdfs for DNA-DNA and peptide-DNA pairs provide clear evidence of two coordination shells. In a control case of dsDNA mixed with a polyK 18-mer, a pronounced, long range order is observed (Fig. S5), which, together with the very low mobility of polyK chains, is in line with the experimental observation of solid precipitates in these systems [28, 33, 37].

Taking advantage of the molecular-scale information provided by the MD simulations, we can discuss the various factors contributing to the onset and stabilization of LC order inside dsDNA coacervates. The fact that NA flexibility tunes the LC stability (Figure 3D) is consistent with previous predictions of theory and simulations [80, 81], as well as the observed decrease of the density at the transition from the LC to isotropic phase. This points to an entropy-driven mechanism of LC appearance, associated with the decrease of excluded volume of the rod-like units upon ordering [82], as often observed for lyotropic LCs. However, the detrimental effect of salt on the stability of the LC phase suggests a role of electrostatic interactions. Indeed, in simulations of PNA-DNA hybrid, which has very similar shape anisotropy to DNA helices, LC ordering is suppressed despite the much higher local concentration. In MD simulations of salt-free coacervates of semi-flexible polyelectrolytes, with a generic CG model lacking any molecular details [81], electrostatic interactions, tuned through the charge of the polycations, were found to facilitate the emergence of LC ordering. According to theoretical studies of salt-free mixtures [79, 80], fluctuation-induced attractive Coulomb correlations between the rodlike polyanions would be responsible for the enhancement of a weakly ordered nematic phase in low-density coacervates. To ascertain the role of peptides in the emergence of LC ordering in our systems, we estimate the average number of contacts, *N*_*c*_, of the amino acids of pL22 with nucleotides inside isotropic and LC coacervates (see SI, Fig. S13). Overall, the pattern of contacts mirrors the charge of amino acids, as reported for proteins [26] and mixtures of peptides with relatively short DNA duplexes [83]. However, for dsDNA *N*_*c*_ is higher in the LC (Fig. S13 B) than in the isotropic phase (Fig. S13 C), which on its turn has a *N*_*c*_ profile close to the isotropic ssDNA (Fig. S13 A). This is consistent with the fact that the two phases, with the same NT/AA ratio (although the ratio of peptides to NA chains in the former is twice as big as in the latter), also display a similar density (see SI, Fig. S5). Interestingly, in the LC phase the interrelated increase of density and onset of orientational order not only bring about an increase of peptide-DNA contacts, but also an increase in the number of DNA-DNA bridges enforced by peptides (see SI, Fig. S14).

In summary, our simulations suggest a crucial and subtle role of electrostatic interactions for the mesomorphic behavior of peptide/dsDNA coacervates, where peptides contribute both through direct interaction with dsDNA, and by mediating DNA-DNA interactions. LC ordering thus entails a strengthening of electrostatic interactions, which in turn stabilizes the ordered phase. If the electrostatic interaction grows too much, as in the case of polyK/dsDNA-20 (see SI, Fig. S13 D) solid precipitates are predicted, in agreement with experiments. We may mention that, within our model, an increase of the ionic strength implies both a decrease of the Debye length and a decrease of the depth of the minimum of the non-bonded pair potential (Eqs. S1 and S3). In real systems, a more complex mechanism may occur, involving ion exchanges and hydration effects.

### Different mobility of oligonucleotides and peptides in condensates

The asymmetry in the structural arrangement between DNA and peptides is reflected in the mobility measured in simulations, which strongly depends on the molecular structure and on the environment [84]. The DNA and peptide concentrations inside simulated droplets are similar to the experimental ones, despite the different preparation concentration (Fig. S3). However, because in CG models the diffusion time scale is accelerated due to the lack of hydrodynamic interactions and molecular details [16, 62, 85], we report scaled diffusion coefficients, taking the value for peptides in the isotropic phase as a reference. Inside isotropic droplets formed by ssDNA, or dsDNA at high ℐ 𝒮, the two species display a similar diffusion coefficient, with peptides slightly more mobile than DNA filaments (Figure 7C, *top row*). On the contrary, in LC droplets of dsDNA, while the dynamics of ordered DNA duplexes is strongly reduced (see also SI, Fig. S15), the peptides surrounding and bridging them keep a similar diffusivity to the isotropic case, with faster motion parallel to DNA chains than perpendicular to them. This suggests that peptide bridges between DNA filaments are highly dynamic [86]. Indeed, also the exchange rate between the condensates and the dilute phase (Figure 7D, where positive and negative values indicate molecules entering and exiting the droplets, respectively) shows a similar pattern for interface mobility, with a fast kinetics of peptides for both isotropic and LC coacervates, while for dsDNA the exchange between LC domains and surnatant is suppressed.

To verify these simulation findings, we experimentally track the mobility of both components inside the condensates by labeling them with different fluorescent dyes and performing Fluorescent Recovery After Photobleaching (FRAP) experiments [87, 88] in the isotropic and LC cases (see SI, Fig. S16). By locally bleaching a portion of the droplet and measuring the evolution of the fluorescent signal in the bleached region, we can separately measure the effective diffusivity of DNA and peptide (Figure 7C, *bottom row*). Similarly to simulations, we find that while the translational diffusion of dsDNA in the LC domains is much slower than that of ssDNA in the isotropic droplets, the mobility of peptides in the two cases is very similar. Finally, the strong mobility reduction of LC dsDNA is also manifested on a larger scale, in coalescence events of coarsening droplets. As shown in confocal microscopy frames in Figure 7E, where ONTs are tagged with a fluorescent dye, the characteristic fusion time, which depends on the viscosity (and surface tension) of the concentrated phase [48, 84], is much faster for ssDNA (top row) than for LC dsDNA (bottom row).

## Conclusions

This study provides a systematic analysis of how NA properties govern peptide-mediated phase separation *via* complex coacervation, with full phase diagram mapping and characterization of the emerging LC ordering. Using a biologically relevant peptide, we disentangle the effects of NA length and secondary structure, establishing robust quantifiers for charge density and flexibility of ONTs. A CG model incorporating hybridization reproduces the experimental phase diagrams across temperature and ionic strength variations, providing molecular level insights. Enhanced LLPS in structured DNA reflects a trade-off between the stabilizing effect of a higher charge density and the opposing influence of stiffness and hidden nucleobases. This subtle balance may underlie conflicting literature reports.

A distinct effect of double stranded DNA is its ability to induce liquid-crystal ordering within droplets, and we show that this occurs in the presence of moderately charged peptides. We find that inside LC droplets, mobile peptides transiently bridge stiff dsDNA tracts, retaining a disordered arrangement and high mobility. These transient bridges, coupled with the entropic gain arising from reduced excluded volume of DNA helices, contribute to the stabilization of the ordered phase. By integrating computational modeling with experimental validation, we establish quantitative tools to link NA sequence and structure to condensate properties. These insights may enable a better design of responsive coacervate systems with applications in synthetic biology, cellular phase separation studies, and nucleic-acid–based biomedical materials.

## Materials and Methods

### Sequence design

#### Tuning secondary structures

Sequence optimization was performed with the aid of the *NUPACK Python module*[89]: given a seed DNA sequence, random permutations (which preserve length and NT composition) are iteratively applied and only the structural ensembles matching the desired criteria are kept, while the others are discarded. The selection criteria are based on: (i) the number of base-paired nucleotides in the *Minimum Free Energy* structure; (ii) the concentration of inter-strand constructs made up of 2 sequences; (iii) the fraction of unpaired nucleotides at equilibrium. For all sequence selections, the default *dna04* model and the *stacking* ensemble were used, and a NUPACK test tube was created containing the seed sequence at the same concentration used in the experiments; moreover, the analysis is performed at *T* = 23^°^C and 100 mM NaCl, mimicking the conditions of LLPS before and after temperature annealing. This approach allowed for the selection of ssDNA-10, ssDNA-20, ssDNA-40 and hp5DNA. Instead, dsDNA-10, dsDNA-20, dsDNA-40 and hdsDNA were obtained by hybridization of the aforementioned ssDNA sequences with their complementary traits (see Table S2), either full or partial.

### Molecular Dynamics simulations

#### Coarse grain model

We used the CG model developed in Refs. [16, 43, 44], where peptides and NAs are described as chains of linked soft beads, each corresponding to an amino acid or a nucleotide, while an implicit solvent representation is used and the effect of counterions and salt is introduced through a Debye-Hückel potential. The model, proposed in Ref. [44] for RNA, was adapted to describe ss- and dsDNA, hybrid structures and polyphosphate (see SI for details).

#### Simulation setup

Phase separation was investigated using the slab method [16]. Initially, straight chains were randomly placed inside a cubic box with periodic boundary conditions in all three directions. Energy minimization was performed to remove steric clashes among the chains. Then, the system was equilibrated in the NPT ensemble (100 *ns*) at a temperature of 150 *K* and a pressure of 1 bar. During this process, the box was compressed until reaching the equilibrium size. Subsequently, the box was elongated in one dimension (*Z* axis) to obtain the desired concentration and the system was brought to the target temperature by a 100 *ns* trajectory in the NVT ensemble. Finally, a 5 *µ*s simulation in the NVT ensemble was carried out, which includes 1 *µ*s of equilibration followed by 4 *µ*s of production. No major changes were observed by increasing the length of the production run.

To build the phase diagrams, several simulations were run at different temperatures and ionic strengths. Samples of ∼5000 beads (around 200 peptides and a number of polyanions so as to neutralize the positive charges) were used, at 120 mM polycation concentration. We performed also a few simulations at the same peptide concentration used in experiments (i.e. *C*_*pL*22_ = 300 *µ*M), taking the same number of polycations and polyanions as in the more concentrated samples and elongating the *Z* axis of the box in the slab method, to reach the target concentration. To check for finite-size effects, for some of the systems additional simulations were performed doubling the size of the sample (∼10000 beads), but no significant changes were observed.

For specific state points, we performed bulk simulations of larger systems (∼ 25000 beads), using periodic boundary conditions in the NPT ensemble with zero external pressure [81]. The simulations started from randomly placed and oriented chains, at a density equal to the density of the clusters obtained by slab simulations at the same temperature and ionic strength.

All simulations were carried out using the LAMMPS package [90]. The equations of motion were integrated with the velocity Verlet algorithm using a timestep of 10 *fs*. Temperature was controlled by a Langevin thermostat [91] with a damping parameter of 100000 *fs*; for NPT simulations, a Nose-Hoover barostat [92] with a damping parameter of 1000 *fs* was applied. Pair interaction potentials were provided in a tabular form. See SI for more details.

#### Analysis of trajectories

Trajectories were analyzed using home-developed *python* scripts and routines from the package *MDAnalysis*[93]. Phase separation was monitored through the distribution of cluster sizes, calculated over the last 4 *µ*s of the trajectory. Phase separation was assumed when the majority of peptides and DNA oligomers (over 50 %) was contained in a large persistent cluster, in a bath of small aggregates, each made of very few chains. The opposite situation, with chains distributed in small clusters, was identified as a single phase. In a few cases, an intermediate behavior was also observed, with one or two relatively big clusters, containing 20 *−* 30 % of polymers, with a short lifetime, together with very small aggregates. For the structural characterization of the condensates we calculated the gyration tensor of the clusters and of the peptide and NA chains, the Radial Distribution Function (RDF) of the beads and the ⟨*P*_2_⟩ order parameter. The mobility of the chains was characterized by the Mean Square Displacement (MSD) of the center of mass of the chains and by the number of exchanges of peptides/DNA with outside the clusters. See SI for more details on these descriptors. To improve sampling, RDFs, translational diffusion coefficients, ⟨*P*_2_⟩ order parameters and average number of contacts were calculated from bulk simulations.

### Quantification of chain flexibility

#### oxDNA simulations

The *oxDNA* CG model [94] (lorenzo-rovigatti.github.io/oxDNA) was employed to perform single-molecule MD simulations of the DNA sequences used in this study. We adopted the sequence-specific *DNA2* parameterization, which accounts for base-specific stacking and hydrogen bonding, and allows for tuning the monovalent salt concentration. Initial conformations were built using *oxView* [95] and subsequently subjected to energy minimization *via* 1000 steps of *Monte Carlo* sampling. Before the actual production simulation, an additional MD relaxation phase of 10^7^ steps using a *Langevin thermostat* with a lowered diffusion coefficient (*D* = 0.5 *SU*) and a small integration time step (*dt* = 0.0005 *SU*) is performed. Starting from the relaxed configuration, the production MD phase in the NVT ensemble (*Andersen thermostat, D* = 2.5 SU) lasted for ∼1.8 *µ*s (*dt* = 0.003 SU). Configurations were printed every 10^4^ steps only after an initial equilibration period of 10^6^ steps. All simulation phases described are performed at 25^°^C and at 50 mM salt concentration. Trajectories were aligned and analyzed with the aid of the *oxDNA Analysis Tools*[95] package.

#### RMSD calculation

The flexibility of DNA oligomers was estimated using a normalized Root Mean Square Deviation (here referred as *RMSD*_*n*_) to account for the intrinsic length dependence of RMSD [76] in short and highly charged systems. To compute the RMSD time series, the centroid structures and configurational ensembles of all oligonucleotides were obtained by *oxDNA* simulations, as described above. For the G-quadruplex sequence in Fig. S11, the NMR ensemble in PDB 1AXV was used and the centroid structure and unweighted deviations were calculated from the 20 configurations depleted from hydrogens and *K*^+^. Instead, the configurations of polyP were sampled by a 100 *ns* long all-atom MD trajectory of a fully deprotonated 20mer in water [96], in the presence of 22 *Na*^+^ ions. Standard parameters of the OPLS-AA force field [97] were used for polyphosphate and *Na*^+^ ions, together with the TIP3P water model. Simulations were performed using GROMACS (www.gromacs.org).

Given a set of configurations of a flexible chain made of *N* units, the RMSD of the *k*-th configuration with respect to a reference structure can be defined as:

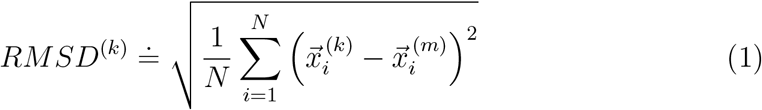

where *N* is the number of monomers and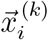 and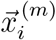 are the coordinates of the *i*-th monomer in the *k*-th configuration and in the reference structure, respectively. In our calculations, we took as a reference the centroid of the structures comprised in the configuration set. Because the time-averaged RMSD scales linearly with the length of the DNA strands (see SI, Fig. S10), we implement the simple normalization:

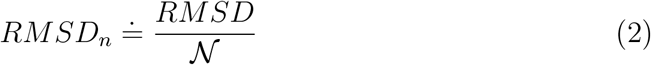

where 𝒩 is the number of monomers of the ONT, i.e., single nucleotides for ssDNA, base pairs for dsDNA, quadruplets for G-quadruplexes, and the proper combination for partially hybridized structures (in analogy with the definition of ℒ _*c*_).

### Calculation of the average surface electrostatic potential

To calculate the average electrostatic potential generated on the surface of a polyanion by its charge distribution, we used atomistic representations of the centroid structures of the oligomers, obtained for each system as described in the previous Section (CG coordinates from *oxDNA* configurations were transformed into PDB using *tacoxDNA*, tacoxdna.sissa.it). Starting from the atomic coordinates, the partial charges and atomic radii, defined according to the AMBER03 force field, were obtained using the APBS-PDB2PQR software suite [98]. The molecular surface was then defined by a finite set of points, determined over a uniform grid in a rectangular box containing the molecule. The Coulomb potential *ϕ*_*e*_ was evaluated at each point, assuming a relative dielectric constant *ϵ* = 2 inside the polyanions. The average surface potential Φ_*e*_ was calculated over the grid by discrete summation:

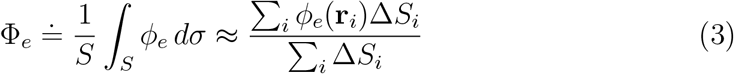

where *S* is the surface area and Δ*S*_*i*_ are surface elements.

The defined Φ_*e*_ bears an intrinsic dependence on the polymer length, arising from the long-range nature of the coulomb potential and from the combinatorial growth of the grid points where the potential is evaluated. Therefore, in a similar fashion to the definition of the *RMSD*_*n*_, we normalize the average surface potential Φ_*e*_ as (see SI for details):

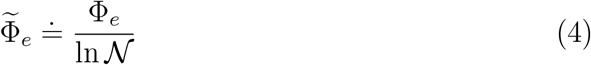

### Materials

#### Oligonucleotides

ssDNA oligomers (table S2) were purchased from IDT (Integrated DNA Technologies) and were used without further purification (standard desalting). Lyophilized oligomers were resuspended in Milli-Q water at 100 −500 *µ*M concentrations and stored at − 20 ^°^C in ∼10 *µ*L aliquots.

Hybridized DNA sequences (Table S2) were obtained by mixing 1:1 the sequence of interest with its complementary segment and by performing a slow temperature annealing with a thermoblock, heating up the solution at 95^°^C for 5 minutes and slowly let it cool down to ambient temperature by thermalisation.

Purified linear polyphosphate at 200 mM concentration was kindly supplied by RegeneTiss.Inc (Japan) in the form of EX-polyP^®^ with average chain lengths of 20. Stock solutions were subsequently diluted down to 500 *µ*M and stored at −20 ^°^C.

HPLC purified PNA (a DNA analog where the negatively charged sugar-phosphate backbone is substituted by a neutral pseudo-peptide skeleton composed of N-(2-aminoethyl)-glycine units) was purchased from Biomers.net GmbH (Table S2) and subsequently dissolved in Milli-Q water at 500 *µ*M concentration. To aid dissolution, the solution was kept shaking at 600 rpm and at 60 ^°^C for several minutes. UV absorbance quantification on a NanoReady Touch micro-volume spectrophotometer of a serial dilution of the PNA stock was in accordance within a 10% error with the nominal concentration, which is satisfactory for the accuracy of such measurements.

The PNA-10 hybrid (Table S2) was obtained by mixing 1:1 the PNA stock solution with its DNA complementary. Before performing a temperature annealing, the solvent conditions were adjusted to 10 mM NaCl, 20 mM Tris-HCl pH 7.7 and 5% *v/v* acetonitrile. The annealing was performed with a thermoblock, heating up the solution at 95 ^°^C for 5 minutes and slowly let it cool down to ambient temperature. The measured UV absorbance melting curve (see SI, Fig. S17) confirmed that hybridization took place (estimated melting temperature of approximately 58 ^°^C).

#### Peptide

pL22 is an ancestral protein fragment of the large ribosomal subunit of *Thermus thermophilus* (see PDB 4V51). It is a basic peptide of 18 AA, enriched in Arg and Lys (table 1) and lacking a well-defined secondary structure in solution, although in the ribosome it adopts a peculiar *β*-loop folding as part of a larger domain.

The pL22 lyophilized stock was synthesized by the Spyder Institute using standard protocols for solid-phase peptide synthesis; it was further dissolved in 20 mM Tris-HCl buffer (pH 7.7) at 2 mM concentration and stored at -20^°^C in ∼10 *µ*L aliquots.

### Determination of phase diagrams

#### Sample preparation

In our study, we consider mixtures with ratio of positive-to-negative polyelectrolyte charges equal to one. Thus, when preparing samples with different ONTs (where the length or the number of charges-per-monomer change), the stoichiometric composition of the mixture changes accordingly, at fixed pL22 concentration (if not stated otherwise, *C*_*pL*22_ = 300 *µ*M).

Stock solutions of NaCl (concentrations of 1 M, 500 mM, 200 mM and 100 mM) and of Tris-HCl (pH 7.7, 100 mM concentration) were prepared, in addition to oligonucleotides and peptide stock solutions. Bovine Serum Albumin (BSA, Sigma-Aldrich) solution was prepared at 3 % mass fraction by dissolving BSA powder in Milli-Q water.

BSA was used to coat plastic well plates (Amine Treated 96 well plates) by adding ∼ 80 *µ*L of solution for about 20 minutes, removing it and washing by adding ∼100 *µL* Milli-Q water for about 10 minutes; this washing step was repeated two times.

Stock solutions were used to prepare coacervate samples of total volume of 10 *µ*L in 200 *µ*L Eppendorf tubes, mixing the components in this given order: Milli-Q water, Tris-HCl buffer, NaCl, pL22 and polyanions. For all the samples, the ionic strength of the mixture is estimated as the molar concentration of added salt plus a 15 mM tare given by the 20 mM Tris-HCl buffer. Mixtures were prepared at room temperature and after the final addition of polyanions the solution was mixed by gentle pipetting. After a few seconds, coacervate samples were transferred in the well plates previously treated with BSA, by locating them in the center of each well. To avoid contamination and evaporation of the samples, each well was covered with ∼ 100 *µ*L mineral oil (M5904, Sigma-Aldrich) and a plastic PCR film was applied on the plastic plate.

#### Optical microscopy

Samples were imaged using a Nikon TE200 microscope, equipped with a Nikon DS-5M CCD camera and temperature-controlled chamber. Images of 2880*×* 2048 pixels were captured at 10×, 20× and 50× magnifications under coherent bright field illumination and typical exposures of 10 *−* 20 ms. To study the phase behavior of coacervate samples as a function of temperature, the following protocol was used. Coacervate droplets were let sediment at 24 ^°^C for about 30 minutes and samples were successively imaged. Samples were then heated up to 90 ^°^C at 1.5 ^°^C*/*min and, after about 10 minutes at 90 ^°^C, were imaged again. Subsequently, samples were cooled down at 1.5 ^°^C*/*min with 5 ^°^C steps and phase state was annotated at each temperature step after a few minutes of equilibration. In some cases, images were captured at every step. At the end of the cooling ramp, samples were imaged at 20 ^°^C and then at 10 ^°^*C* after a final cooling step at 1.5 ^°^C*/*min. All temperature-salt phase diagrams reported in the main text refer to data acquired during the cooling ramp.

### Raman Microscopy

Raman spectra were acquired using a Raman microspectrometer (InVia Reflex, Renishaw plc, Wotton-under-Edge, UK) equipped with a 532 nm laser line. Raman measurements on coacervates were performed using an excitation laser at 100% power (50 mW nominal value at the source), a 1800 L/mm grating and a Leica 100× objective; for each sample, a 10 *µ*L aliquot was sandwiched through a Raman grade *CaF*_2_ disc and a glass cover slip, separated by 50 *µ*m spacers.

Spectra from at least five distinct coacervates were acquired, each spectrum recorded as a sum of 5 accumulations of 2 s each. Coacervate Raman imaging was performed under the same conditions, with spectra collected every 0.5 *µ*m and an accumulation time of 0.5 s per point. For the measurement of dried reference samples, a 5 *µ*L drop of molecules dispersed in water was deposited on the surface of a Raman grade *CaF*_2_ disk and left drying under flow for 20 minutes. Spectra were acquired using the same instrument and settings, and each spectrum was recorded as the sum of ten 3 s accumulations.

Spectra were pre-processed using the *RamanSPy python* package [99]. For both dry and coacervate spectra, a custom pipeline is built to perform: (i) cropping in the spectral region [600, 1800] cm^*−*1^ for dry samples or [1150, 1750] cm^*−*1^ for coacervate samples; (ii) cosmic rays removal using the Whitaker-Hayes algorithm (default parameters); (iii) spectral denoising using a Savitzky-Golay filter (window = 15 and polynomial order = 2); (iv) baseline correction based on asymmetric least squares (smoothing constant = 1000000, differential order = 2 and max iterations = 50); (v) vector normalization. Spectra were then averaged and deviations computed as standard errors.

Raman spatial maps were obtained by scanning a region of interest comprising of one or more condensates and a supernatant portion. Raman spectra were acquired at each point with similar settings as the ones used for point measurements. For pre-processing and analysis of the maps, the Orange software (version 3.36.1) was used and the following pipeline is implemented: (i) cropping in the spectral region [1150, 1750] cm^*−*1^; (ii) rubber band baseline correction; (iii) spectral de-noising using a Savitzky-Golay filter (window = 11 and polynomial order = 2), followed by gaussian smoothing (one standard deviation) and PCA denoising with 4 components; (iv) vector normalization.

### Mobility

Fluorescently labeled ssDNA-20 and dsDNA-20 coacervate samples were prepared for confocal microscopy according to the procedure for phase diagram characterization, but with the addition of TAMRA-labeled pL22 and FAM-labeled DNA added in a 1:200 ratio with respect to the unlabeled molecule concentrations.

Immediately after preparation, samples were sandwiched between two glass BSA-coated coverslips, separated by a silicone gasket of thickness 1.0 mm with a circular aperture of diameter 9 mm (GraceBio-Labs FastWells™ reagent barriers). After the cell was sealed, the samples were then allowed to equilibrate for about 2 hours at room temperature.

### FRAP measurements

The liquidity of the coacervate samples described in the above Section and the molecular mobility within the condensates were assessed by Fluorescence Recovery After Photobleaching (FRAP). FRAP measurements were performed exploiting a laser scanning Confocal Microscope (Leica Stellaris 8) equipped with a 63*×* (*NA* = 1.40) oil immersion objective (HC PL APO CS2) and a sCMOS camera Leica DFC9000 GTC.The imaging system is operated using the LASX (LEICA) Software.

Photobleaching is achieved by using a 495 nm laser at maximum output for 2-5 cycles for the FAM fluorophore or a 530 nm laser at maximum output for 5-10 cycles for TAMRA (each cycle lasting approximately 500 ms). For a given droplet, bleaching is performed on a circular region of interest (ROI) of diameter of the order of 1 *µ*m. For each bleached ROI, a sequence of 5 frames is recorded prior to bleaching (typical duration of 5 s) and one of 100-200 frames immediately after bleaching (typical duration of 1-2 minutes). To monitor photobleaching during the acquisition, only droplets close to at least one other droplet, chosen as a reference, are considered.

For each FRAP measurement, the pre- and post-bleaching raw data series are analyzed without further image pre-processing. Using Image-J, a circular ROI corresponding to the bleach spot is drawn on the first frame of the post-bleaching series and the average intensity *I*_*post*_(*t*) is measured; using the same exact ROI, the pre-bleaching average intensity *I*_*pre*_(*t*) is extracted. Moreover, a round ROI of variable size is drawn outside any droplet to quantify the average background intensity *I*_*bkg*_(*t*), while onw is drawn on a reference unbleached droplet to estimate the unwanted photobleaching *I*_*blch*_(*t*).

The *I*_*pre*_(*t*), *I*_*post*_(*t*) and *I*_*blch*_(*t*) signals are first corrected by subtracting the background intensity 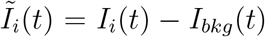. Then, the post-bleaching intensity is normalized with respect to the unwanted photobleaching, 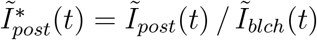, and the same is done for the time-averaged pre-bleaching intensity, 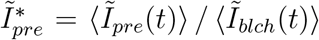. The normalized integrated time-dependent concentration after photobleaching is then defined as:

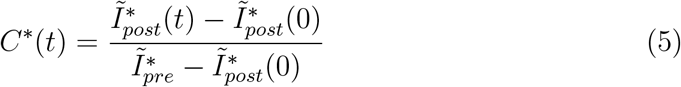

The time evolution of *C*^*∗*^(*t*) is modeled with a simple exponential recovery:

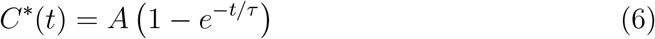

where *A* is the fractional recovery and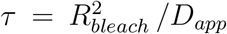, with *D*_*app*_ being the apparent diffusion coefficient. Fitting the above equation to the experimental concentration profile allows for the estimation of *D*_*app*_ and *A* as the best-fitting parameters.

## Abbreviations

AA: (amino acid)
bp: (base pair)
CG: (coarse grain)
ds: (double-stranded)
FENE: (finitely extensible non linear elastic)
LC: (liquid crystal)
LLCPS: (liquid-liquid crystal phase separation)
LLPS: (liquid-liquid phase separation)
MD: (molecular dynamics)
MSD: (mean square displacement)
NA: (nucleic acid)
NT: (nucleotide)
ONT: (oligonucleotide)
polyK: (poly-L-lysine)
PNA: (peptide nucleic acid)
polyP: (polyphosphate)
rdf: (radial distribution function)
ss: (single-stranded)

## Author Contributions

D.A. performed and analyzed the simulations, S. C. performed and analyzed the experiments, A. F. and G. Z. designed and supervised the research, C. M. performed Raman imaging. All authors contributed to interpreting and presenting the results and to writing the paper, and they all reviewed the manuscript.

## Acknowledgments

The authors thank Tommaso Bellini, Tommaso Fraccia and Marco Todisco for useful discussions. We gratefully acknowledge the gift of polyP by Toshikazu Shiba (Regenetiss Inc., Japan). A.F. and D.A. acknowledge funding from the European Union - Next-Generation EU (PNRR M4C2-Investimento 1.4-CN00000041) CN3 Spoke 7 - CUP C93C22002780006 - National Center for Gene Therapy and Drugs based on RNA Technology. G.Z. and S.C. acknowledge funding from SEED4EU+ RECHARGE. We acknowledge the C3P HPC facility at the Department of Chemical Sciences of the University of Padova, the CAPRI initiative (Calcolo ad Alte Prestazioni per la Ricerca e l’Innovazione, University of Padova Strategic Research Infrastructure Grant 2017) and the CINECA grants (HP10CS3JKT, HP10C5KO39) under the ISCRA initiative, for the availability of high-performance computing resources and support. The use of a confocal microscope was supported by the Department of Excellence Project ‘SCALE UP’ funded by the Italian Ministry of University and Research (MUR), Department of Medical Biotechnology and Translational Medicine.

## Supplemental Information

Additional supplemental information can be found online in the Supplemental Information section.

