## Supplemental information for "The Dark Side of Biomolecular Condensates: Quantifying the Role of Nucleic Acids"

[1]Department of Chemical Sciences, University of Padova, Via Marzolo 1, 35131 Padova, Italy
[2]Department of Medical Biotechnology and Translational Medicine, Università degli Studi di Milano, via F.lli Cervi 93, 20054 Segrate (MI), Italy
[3]Laboratory of Nanomedicine, Istituti Clinici Scientifici Maugeri IRCCS, Via Maugeri 10, 27100 Pavia, Italy
[4]These authors contributed equally

### Molecular Dynamics simulations

**Coarse grain model** We used the CG model developed in Refs. [1–3], where peptides and NAs are represented as chains of linked soft beads, each corresponding to an amino acid or a nucleotide. Here we extended the model to describe fully and partially double-stranded structures, including hybrid PNA-DNA duplexes, as well as polyphosphate chains. The interaction potential between a pair of beads is expressed as the sum of bonded and non-bonded terms [1].

The non-bonded potential contains a short-range ($U_{\mathrm{HPS}}$) term and, for NTs and charged amino acids, an electrostatic contribution ($U_{\mathrm{DH}}$). The short-range potential has the form proposed by Ashbaugh and Hatch [4] to describe the effective interaction between amino acids:

$$U_{\mathrm{HPS}}(r_{ij}) = \begin{cases} U_{\mathrm{LJ}} + (1-\lambda_{ij})\epsilon & \text{if } r_{ij} \leq 2^{(1/6)}\sigma_{ij} \\ U_{\mathrm{LJ}}\lambda_{ij} & \text{otherwise} \end{cases} \tag{S1}$$

where $r_{ij}$ is the distance between the beads $i$ and $j$ and $U_{\mathrm{LJ}}$ is the pure Lennard-Jones potential

$$U_{\mathrm{LJ}}(r_{ij}) = 4\epsilon\left[\left(\frac{\sigma_{ij}}{r_{ij}}\right)^{12} - \left(\frac{\sigma_{ij}}{r_{ij}}\right)^{6}\right] \tag{S2}$$

with $\epsilon$ a quantity that determines the absolute energy scale of the short-range interactions and $\sigma_{ij}$ related to the diameters of the beads. The hydropathy parameter $\lambda_{ij}$ in Equation (S1) is introduced to tune the attractive force between beads according to their hydrophobicity. At $r_{ij} = 2^{(1/6)}\sigma$ the potential takes the value $U_{\mathrm{HPS}} = -\epsilon\lambda_{ij}$ and the interaction force vanishes. At shorter distances the force is repulsive and is identical for all beads; at longer distance the force is attractive for $\lambda_{ij} > 0$ (hydrophobic interaction), whereas it remains repulsive for $\lambda_{ij} \leq 0$ (hydrophilic interaction).

The electrostatic interaction is modeled by the Debye-Hückel potential:

$$U_{\mathrm{DH}}(r_{ij}) = \frac{Z_i Z_j e^2}{4\pi\epsilon_0\epsilon_r r_{ij}} e^{-\frac{r_{ij}}{\lambda_{\mathrm{D}}}} \tag{S3}$$

where $Z_i$ and $Z_j$ are the charge numbers of the beads, $\epsilon_0$ is the vacuum permittivity, $\epsilon_r$ is the (relative) dielectric constant of water, $e$ is the elementary charge, and $\lambda_D$ is the Debye screening length:

$$\lambda_{\mathrm{D}} = \sqrt{\frac{\epsilon_0\epsilon_{\mathrm{r}} k_{\mathrm{B}} T}{2000 N_{\mathrm{Av}} e^2 \mathcal{I}S}} \tag{S4}$$

with $N_{\mathrm{Av}}$ the Avogadro number, $k_{\mathrm{B}}$ is the Boltzmann constant, $T$ is the temperature, and $\mathcal{I}S$ (in mol/L) is the ionic strength.

The bonded interaction between two beads is described by a harmonic potential [1]:

$$U_{\mathrm{str}}(r_{ij}) = \frac{k_{\mathrm{str}}}{2}(r_{ij} - r_0)^2 \tag{S5}$$

where $k_{\mathrm{str}}$ is the spring constant and $r_{ij,0}$ is a reference distance.

To model base-paired tracts, we introduced two additional terms. A stretching potential in the form of Equation (S5) with constant $k_{\mathrm{bp}}$ was used to link beads representing H-bonded pairs. Moreover, a harmonic bending potential between three consecutive beads in a strand of base-paired nucleotides was introduced to account for the stiffness of double-stranded tracts:

$$U_{\mathrm{bend}}(r) = \frac{k_{\mathrm{b}}}{2}(\theta - \theta_0)^2 \tag{S6}$$

where $k_{\mathrm{b}}$ is the strength constant and $\theta_0$ is a reference angle between the three beads.

**Model parameters** The specific parameters for amino acids and nucleotides entering Equations (S1)-(S6) were taken from Refs. [2] and [3]. In particular, for the short-range non-bonded potential of amino acids we used values based on the Urry hydropathy scale, derived from experimental data on heat-induced conformational compaction of host-guest polypentapeptides, which were found to provide a good description of the phase separation of disordered proteins [3]. For nucleotides, we used the hydropathy scale proposed by Kapcha and Rossky (KR scale) [5] for proteins, that is based on the atomic charges in the OPLS all-atom force field [6]. The hydropathy parameters for DNA nucleotides were calculated according to the procedure outlined in Ref. [2] for RNA. For adenosine, cytosine and guanine, replacement of an OH by a hydrogen atom in ribose yields slightly larger hydrophobicity parameters; for thymine there is an additional contribution due to replacement of a methyl group in uracil by a hydrogen atom. The parameters of the $U_{HPS}$ potential for all beads are listed in Table 1, along with the charge numbers $Z_i$. The parameter $\epsilon$ in Equation (S1) was set to be constant across all pairs and equal to 0.2 kcal/mol [2], whereas the values of $\sigma_{ij}$ and $\lambda_{ij}$ for beads of different types were computed as the arithmetic means $\sigma_{ij} = (\sigma_i + \sigma_j)/2$ and $\lambda_{ij} = (\lambda_i + \lambda_j)/2$, where a single subscript is used for interactions between beads of the same type. For the dielectric constant $\epsilon$ of water in Equation (S4), temperature dependent values were used [7], ranging from 87.7 (at $T$=273.15 K) to 55.7 (at $T$= 363.15 K) [8].

The total non-bonded potential was truncated at a cutoff distance $r_c$ and shifted in such a way that $U_{\mathrm{HPS}}(r_c) + U_{\mathrm{DH}}(r_c) = 0$. The value of $r_{\mathrm{c}}$ was taken equal to the distance at which the non-bonded interaction potential between two adenine beads, which have the strongest interaction, is equal to 0.005 kcal/mol, in line with Ref. 2. For both peptides and DNA filaments, a stretching constant $k_{\mathrm{str}}$ =2.4 kcal/(mol Å$^2$) was used in Equation (S5), together with a reference distance $r_0$ equal to 3.8 Å for amino acids and to 5 Å for NTs [2, 3]. Moreover, for the stretching potential between base-paired NTs we used $k_{\mathrm{bp}} = 40$ kcal/(mol Å$^2$), with a center-to-center distance equal to the arithmetic mean of their diameters, which is comparable to the average distances between the centers of mass of the NTs in B-DNA [9]. In the bending potential, Equation (S6), considering that the persistence length of dsDNA ($\sim$ 50 nm at 10 mM Na$^+$) is over 50 times greater than that of ssDNA ($\sim$ 0.75 nm at 10 mM Na$^+$) [10], we assumed $k_{\mathrm{b}}$ =40 kcal/(mol rad$^2$) and $\theta_0 = \pi$. To check the effect of flexibility, additional simulations were performed using a bending constant $k_{\mathrm{b}} = 2$ kcal/(mol rad$^2$) for ssDNA and for dsDNA.

For PNA units, we used the same model and the same parameters as for DNA, with the only difference that charges were switched off. PolyP was modeled as a chain of soft beads with a reference distance of 3 Å, a diameter equal to 6 Å, a charge of -1 $e$ and a hydropathy parameter equal to 0. A constant $k_{\mathbf{str}} = 1$

kcal/(mol Å$^2$) in the stretching potential, Equation (S5), was assumed, which yields a persistence length, calculated from the decay of the autocorrelation function of bond vectors, similar to that of ssDNA.

| AA/NT | $Z_i$ | $\sigma_i$ (Å) | $\lambda_i$ |
|---|---|---|---|
| ALA | 0.0 | 5.04 | 0.60 |
| ARG | 1.0 | 6.56 | 0.56 |
| ASN | 0.0 | 5.68 | 0.59 |
| ASP | -1.0 | 5.58 | 0.29 |
| CYS | 0.0 | 5.48 | 0.65 |
| GLN | 0.0 | 6.02 | 0.56 |
| GLU | -1.0 | 5.92 | 0.00 |
| GLY | 0.0 | 4.5 | 0.57 |
| HIS | 0.5 | 6.08 | 0.76 |
| ILE | 0.0 | 6.18 | 0.71 |
| LEU | 0.0 | 6.18 | 0.72 |
| LYS | 1.0 | 6.36 | 0.38 |
| MET | 0.0 | 6.18 | 0.68 |
| PHE | 0.0 | 6.36 | 0.82 |
| PRO | 0.0 | 5.56 | 0.76 |
| SER | 0.0 | 5.18 | 0.59 |
| THR | 0.0 | 5.62 | 0.59 |
| TRP | 0.0 | 6.78 | 1.00 |
| TYR | 0.0 | 6.46 | 0.90 |
| VAL | 0.0 | 5.86 | 0.66 |
| ADE | -1.0 | 8.44 | 0.08 |
| CYT | -1.0 | 8.22 | 0.11 |
| GUA | -1.0 | 8.51 | -0.05 |
| THY | -1.0 | 8.17 | 0.19 |

Table S 1: Charge numbers and parameters of the non-bonded short-range potential $U_{PS}$ for amino acids and nucleotides, used in the CG-MD simulations.

**Analysis of trajectories** The radial distribution function, RDF, was calculated using the routine available in MDAnalysis.[11] The other quantities used to characterize structural (gyration tensor, $\langle P_2 \rangle$ order parameter, average number of contacts of peptide residues with NTs) and dynamic properties (mean square displacement and translational diffusion coefficient, number of chain exchanges in/out of clusters) were calculated using home-made codes.

**Gyration tensor** The gyration tensor of a chain or of a cluster was calculated as

$$\mathbf{T}_{\mathrm{g}} = \frac{1}{N_{\mathrm{b}}} \sum_{i=1}^{N_b} (\vec{r}_i \otimes \vec{r}_i - \vec{r}_{\mathrm{CM}} \otimes \vec{r}_{\mathrm{CM}}) \tag{S7}$$

where $N_{\mathrm{b}}$ is the number of beads, $\vec{r}_i$ is the vector position of the $i$th bead and $\vec{r}_{\mathrm{CM}}$ is the vector position of the center of mass of the chain/cluster.

**Orientational order parameter** The orientational order was quantified through the order parameter $\langle P_2 \rangle$, that is the average value of the second Legendre polynomial, having as argument the cosine OF the angle between the director $\hat{n}$, i.e. a unit vector parallel to the $C_\infty$ symmetry axis of the system, and the long axes of the aligning units. Considering that in our systems the orientational order is determined by the dsDNA chains, the director was calculated by diagonalizing the quadrupolar ordering tensor: [12]

$$\mathbf{Q} = \frac{1}{2N_{\mathrm{DNA}}} \sum_{i=1}^{N_{\mathrm{DNA}}} (3\, \hat{u}_i \otimes \hat{u}_i - \mathbf{I}_3) \tag{S8}$$

where $N_{\mathrm{DNA}}$ is the number of DNA chains, $\mathbf{I}_3$ is the identity matrix, and $\hat{u}_i$ is a unit vector parallel to the alignment axis of the $i$-th chain, which was taken parallel to the major eigenvector of the gyration tensor of the chain. The director $\hat{n}$ is the major eigenvector of the $\mathbf{Q}$ tensor and the corresponding eigenvalue is the order parameter of the DNA chains, $\langle P_2 \rangle_{\mathrm{DNA}}$. The phase was labeled as LC if $\langle P_2 \rangle_{\mathrm{DNA}} \gtrsim 0.8$.

For peptides, we took the chain bonds as the aligning units; thus the order parameter was calculated as:

$$\langle P_2 \rangle_{\mathrm{pep}} = \frac{1}{2N} \sum_{i=1}^{N} (3\, \hat{u}_i \cdot \hat{n} - 1) \tag{S9}$$

where $N$ is equal to the product of the number of peptide chains times the number of bonds in a chain and $\hat{\mathbf{u}}_i$ is a unit vector parallel to a peptide chain bond.

**Definition of clusters and calculation of chain densities** For the definition of clusters we used a bead-to-bead distance metric, i.e., a chain belongs to a cluster if it has at least one bead within a distance $d_c$ from any other chain of the cluster. Based on the radial distribution function calculated for the various species, $d_{\mathrm{c}} = 15$ Å was assumed.

In the presence of phase separation, the dense phase concentration was calculated as the ratio between the mass of the chains contained in the largest cluster and the volume of the cluster. This was estimated as the volume of a sphere with a radius equal to the average gyration radius of the cluster. The density of the dilute phase was then computed from the mass of the remaining polymer chains and from the volume of the box outside the cluster.

**Average number of contacts** To calculate the average number of contacts of amino acids with nucleotides, we counted the NT beads having their centers within a cut-off radius ($r_c$) from the centers of AA beads. Based on the values of $\sigma_i$ (Table 1), $r_c = 9$ Å was assumed.

**Dynamic quantities** The translational diffusion coefficients of chains were determined by linear fitting of the mean square displacements (MSDs) of the centers of mass of single chains *vs* lag time in the diffusion regime. Analogously, for liquid-crystalline systems, we calculated the diffusion coefficients along the director and perpendicular to it. For the calculation of the number of exchanges in/out of a cluster, a chain was assumed to be in the cluster unless all its beads were outside.

### Normalization of the electrostatic potential

For a linear straight polymer with $\mathcal{N}$ charged monomers, each spaced by $l$, a probe charge at distance $x$ experiences a Coulomb potential $A/x$, where $A = e/4\pi\epsilon$. Thus, if the probe is at a distance $d$ from each bead, it will feel an overall potential given by the sum of the potentials for each bead. Without loss of generality, if we consider the case $\mathcal{N}$ odd, the total potential $\mathcal{V}$ can be expressed as the sum of 4 terms:

$$\mathcal{V} = \frac{\mathcal{N}A}{d} + 2\sum_{i=1}^{\frac{\mathcal{N}-1}{2}} \frac{A}{l} + 2\sum_{i=1}^{\frac{\mathcal{N}-1}{2}} \frac{2A}{il}\left(\frac{\mathcal{N}}{2} - i\right) + 2\sum_{i=1}^{\frac{\mathcal{N}-1}{2}} \frac{A}{il} \tag{S10}$$

which (upon proper rearrangement of the terms and evaluation of the identity summations) in the limit $\mathcal{N} \to \infty$ can be approximated by:

$$\frac{\mathcal{V}\,l}{A\mathcal{N}} \simeq \alpha + 2\,(\gamma - \ln 2) - 1 + 2\,\ln\mathcal{N} + 2\,\frac{\ln\mathcal{N}}{\mathcal{N}} + \mathcal{O}\left(\frac{1}{\mathcal{N}}\right) \tag{S11}$$

with $\alpha \doteq l/d$ and $\gamma$ the Euler-Mascheroni constant. Thus the scaling of the average surface electrostatic potential $\mathcal{V}/\mathcal{N} \sim \ln\mathcal{N}$ and the normalization in Equation 4.

### Experimental methods

**Determination of local concentration** To estimate the DNA concentration in the dense phase ($C_d$) for a given stoichiometry ($C_s$), the volume fraction of the dense phase ($\varphi$) and the concentration of DNA in the supernatant ($C_l$) were measured; $C_d$ can be then extracted from mass conservation:

$$C_d = \frac{C_s - C_l}{\varphi} - C_l \tag{S12}$$

To estimate the volume fraction of the dense phase, fluorescently labeled samples were prepared and imaged as described in Materials and Methods, Mobility. A z-stack with equal spacing between slices ($z_{gap} \sim 300$ nm) of a large field of view ($\sim 34116\,\mu$m$^2$) of the sample was acquired. After thresholding of the whole stack using the ImageJ software, the volume fraction $\varphi$ is simply computed as:

$$\varphi = \frac{z_{gap}}{h} \sum_{s} A_f^{(s)} \tag{S13}$$

where $h = 1000\,\mu$m is the height of the sandwiched sample and $A_f^{(s)}$ is the area fraction of pixels belonging to the coacervate phase in the slice $s$.

To estimate the DNA supernatant concentrations, samples were prepared in small Eppendorf tubes and droplets were let sediment at ambient temperature for at least one hour; subsequently, the Eppendorf tubes were centrifuged for 1 minute and few microliters were carefully removed from the supernatant phase. DNA concentrations were then measured *via* standard UV-absorbance at 260 nm, $C_l = A_{260}/(\epsilon_{260}\, l)$, where $l$ is the variable optical path of the micro-spectrometer and $\epsilon_{260}$ is the sequence-specific extinction coefficient of the DNA sequence.

Measurements of both $\varphi$ and $C_l$ were performed in triplicates and the error on $C_d$ was estimated from the error propagation of Equation (S12), where: (i) the error on $C_s$ is computed from the pipetting error due to stock solutions dilutions; (ii) the error on $C_l$ is given from the standard deviation of 3 independent UV-absorbance measurements; (iii) the error on $\varphi$ is given from the standard deviation of 3 z-stacks taken from different field of views.

Table S 2: All the DNA/PNA oligomers employed in this study: abbreviations, primary structures and secondary structures in dot-bracket notation; the & sign indicates ligation of two strands, while bold sequences refer to PNA traits.

| **Abbreviation** | **Primary structure** | **Secondary structure** |
|---|---|---|
| ssPNA | 5'-**GGACGACTTG**-3' | None |
| ssDNA-10 | 5'-GGACGACTTG-3' | None |
| ssDNA-20 | 5'-GTTAAAGTGCCAAGTCGTCC-3' | None |
| ssDNA-40 | 5'-GCATATCATCGGTACACATCATCTCGGCAGGGGTCAGTTA-3' | None |
| ssDNA-80 | 5'-CACAACACCCACAACCAAAAACAACAAACCACACCCCAAC<br>CCACACAAACACACCCCAACAACCACCACAACCAACAACC-3' | None |
| hdsDNA | 5'-GTTAAAGTGCCAAGTCGTCC-3'<br>5'-GGACGACTTG-3' | ..........((((((((((&)))))))))) |
| hp5DNA | 5'-TGCGAGAGATATCTCGATCC-3' | ..(((((....))))).... |
| dsDNA-10 | 5'-GGACGACTTG-3'<br>5'-CAAGTCGTCC-3' | ((((((((((&)))))))))) |
| dsDNA-20 | 5'-GTTAAAGTGCCAAGTCGTCC-3'<br>5'-GGACGACTTGGCACTTTAAC-3' | ((((((((((((((((((((&)))))))))))))))))))) |
| dsDNA-40 | 5'-GCATATCATCGGTACACATCATCTCGGCAGGGGTCAGTTA-3'<br>5'-TAACTGACCCCTGCCGAGATGATGTGTACCGATGATATGC-3' | ((((((((((((((((((((((((((((((((((((((((&<br>&)))))))))))))))))))))))))))))))))))))))) |
| PNA-10 hybrid | 5'-**GGACGACTTG**-3'<br>5'-CAAGTCGTCC-3' | ((((((((((&)))))))))) |

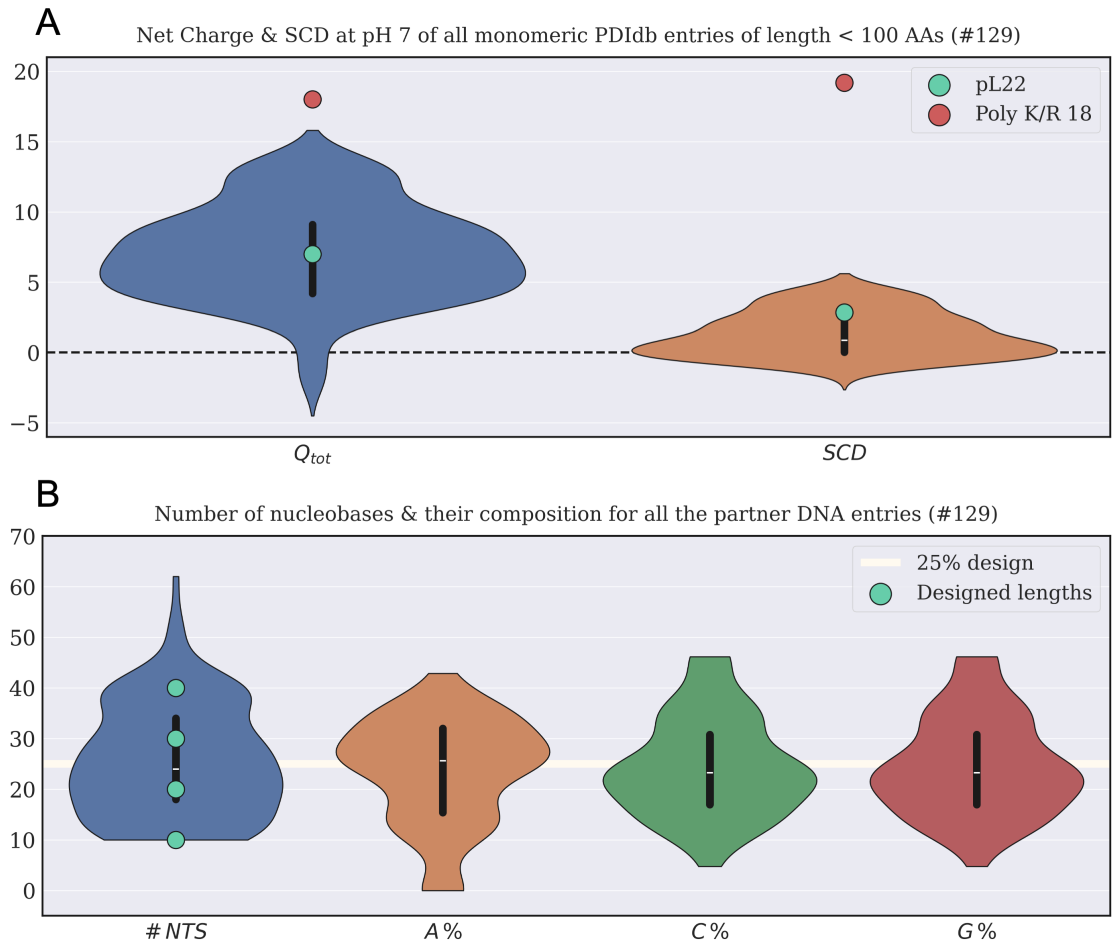


Figure S 1: *Comparison between the pL22 peptide, the designed DNA sequences and a subset of 129 entries from the* PDIdb *repository (http://melolab.org/pdidb/web/content/home), containing relevant structural information of protein-DNA complexes. The selected entries correspond to all the DNA-binding proteins with less than 100 AAs and lacking quaternary structure. A) Violin plots of net charge and Sequence Charge Decoration (SCD) of the proteins at pH 7.0 (R,K = +1; D,E = -1; H = +0.1). The green markers refer to the net charge and SCD of pL22, while the red ones to the same quantities for a Poly-L-lysine or Poly-L-arginine of the same length. B) Violin plots of the number of nucleotides and nucleobases composition in the DNA sequences corresponding to panel A; green markers show the probed DNA lengths in this study.*

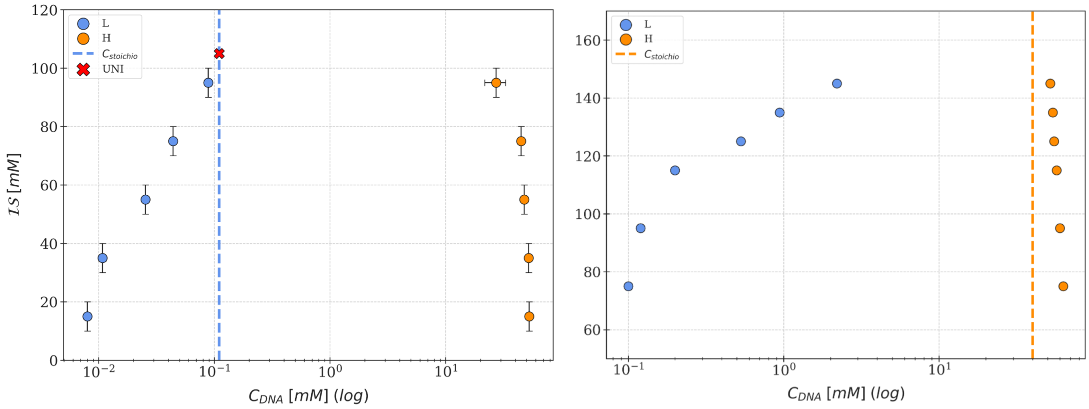


Figure S 2: *Experimental and simulated (marked by "SIM") phase diagrams of mixtures of pL22 with ssDNA-40 (first row, left & right ), ssDNA-80 (second row, left & right), dsDNA-10 (third row, left) and dsDNA-40 (third row, right). For ssDNA-40 the full* $T - \mathcal{IS}$ *phase diagrams are shown, while for the other oligonucleotides the phase behavior was annotated only at* $T = 20\,^{\circ}C$ *(or* $T = 30\,^{\circ}C$ *for simulated ssDNA-80) after temperature annealing. Phases are denoted by the color and shape of markers and shaded regions: monophasic (orange pentagons), isotropic biphasic (green full circles) and liquid-crystalline biphasic (blue diamonds).*

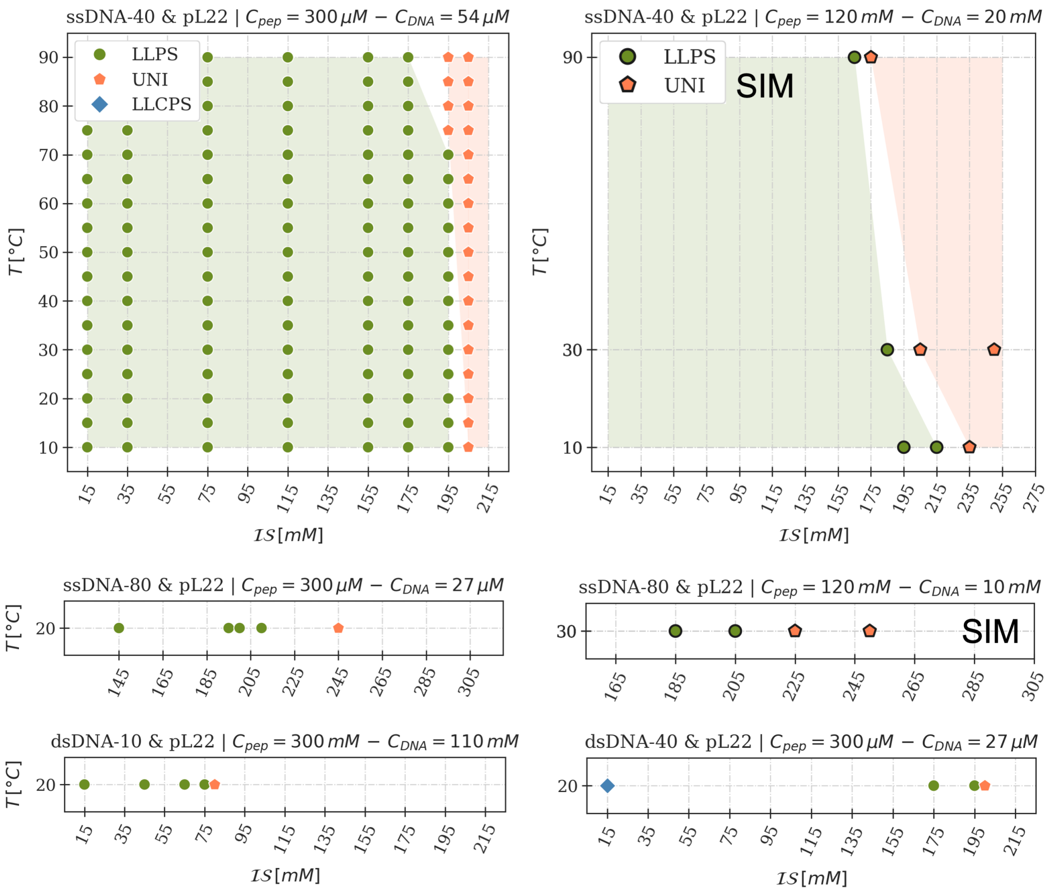


Figure S 3: *Experimental (left) and simulated (right) phase diagrams in the DNA concentration-ionic strength of the solution plane (x-axis, log scale, and y-axis, lin scale, respectively). "UNI" denotes no demixing, while "L" and "R" refer to the "left" and "right" branches of the bell-shaped diagram. Concentrations of the dense phase (R) are estimated from the concentrations of the supernatant phase (L), scanning the line at constant concentration "$c_{stoichio}$" of DNA and pL22: for every given ionic strength, a tie-line connects the concentration of the two phases.*

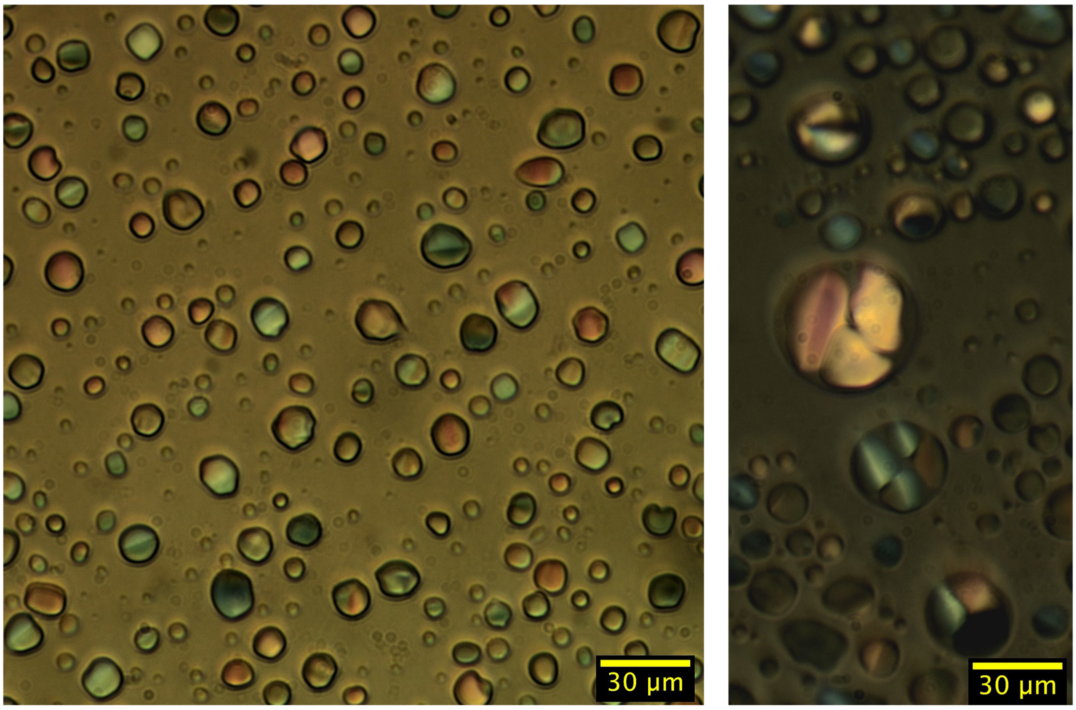


Figure S 4: *Polarized optical microscopy images of dsDNA-20 and pL22 coacervates ($C_{DNA}$=55 μM & $C_{pL22}$=300 μM) at ambient temperature and $\mathcal{IS}$=95 mM (left) or $\mathcal{IS}$=15 mM (right). In both cases, birefringent textures indicate liquid-crystalline ordering.*

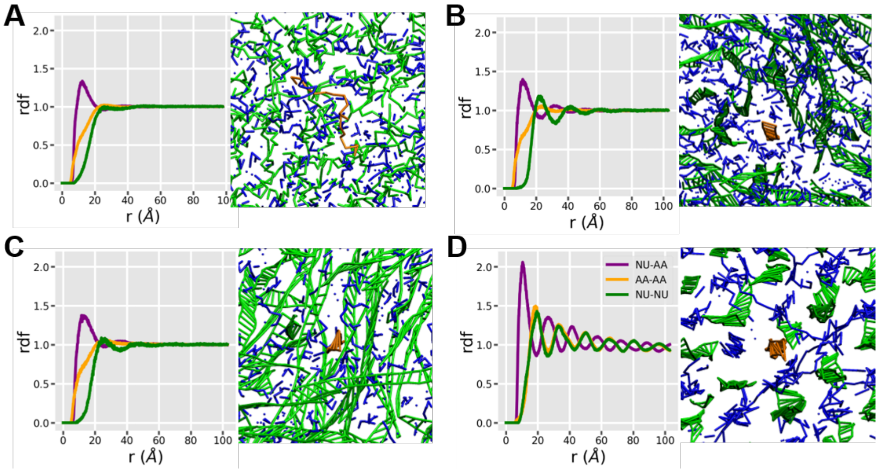


Figure S 5: *Radial Distribution Functions (rdf) inside coacervates (left) and corresponding illustrative snapshots (right) from bulk MD simulations at* $T = 303$ *K of: A) pL22/ssDNA-20,* $\mathcal{IS}$*=75 mM (isotropic); B) pL22/dsDNA-20,* $\mathcal{IS}$*=75 mM (LC); C) pL22/dsDNA-20,* $\mathcal{IS}$*=225 mM (isotropic); D) polyK-18/dsDNA* $\mathcal{IS}$*=75 mM (solid). The plots show the rdf calculated for amino acids belonging to different peptides (orange), for DNA nucleotides belonging to different ONTs (green), and for amino acids around DNA nucleotides (violet). In the snapshots, DNA chains are colored green, whereas blue is used for amino acids within 9 Å from DNA nucleotides; the DNA chain in the middle is highlighted in orange for reference. The rdfs in isotropic coacervates in C are only slightly more structured than those in A, both indicating direct contacts between amino acids and nucleotides, but no contacts between the nucleotides and scarce, yet not fully absent, contacts between peptides. The same holds for LC coacervates in B, but in this case the structure of the rdfs for DNA-DNA and peptide-DNA pairs provides clear evidence of two coordination shells. In D the order is even more pronounced and extends to longer range; moreover, there are no contacts between peptides, which can be attributed to their mutual electrostatic repulsion in the highly charged polyK.*

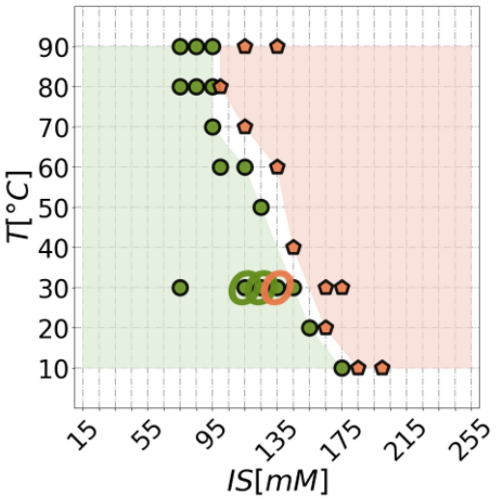


Figure S 6: *Phase diagram of pL22/ssDNA-20 mixtures as a function of the temperature $T$ and of the ionic strength $IS$, obtained from slab-MD simulations starting from $C_{DNA}$ = 40 mM and $C_{pL22}$ = 120 mM. The phase behavior is denoted by the color and shape of markers and shaded regions: monophasic (orange pentagons) and isotropic biphasic (green full circles). The orange/green empty circles show the results obtained using the same concentration used in experiments, namely $C_{DNA}^{sim}$=110 $\mu M$ and $C_{pL22}^{sim}$=300 $\mu M$.*

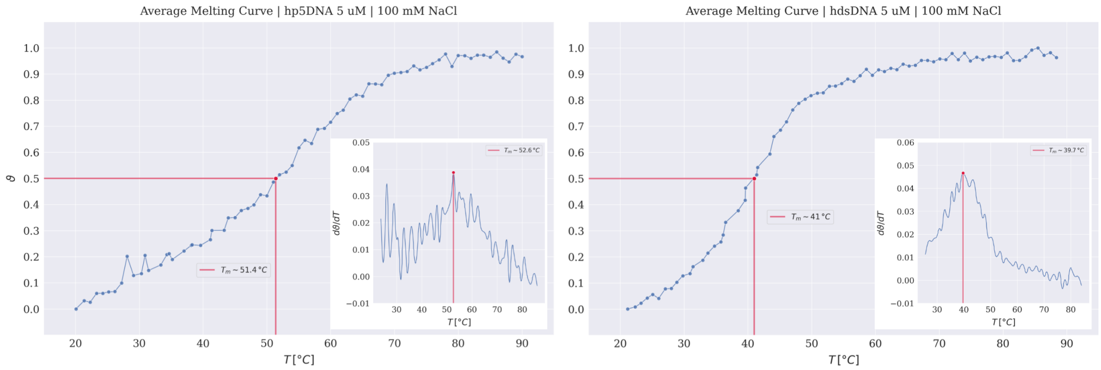


Figure S 7: *Normalized absorbance melting curves of hp5DNA (left) and hdsDNA (right) at 5 $\mu$M in aqueous 100 mM NaCl solution. Absorbance spectra in the 210-320 nm range are collected and the raw signals around the 260 nm peak are then averaged and normalized. The value $\vartheta$=0.5 determines a first estimate of the melting temperature (indicated by the red markers and lines), around 51 °C for hp5DNA and 41 °C for hdsDNA. The insets of the two curves show the numerical differentiation of the melting curves, upon smoothing and interpolation, with the maxima providing a more reliable estimate of the melting temperatures (53 °C for hp5DNA and 40 °C for hdsDNA) due to baseline removal.*

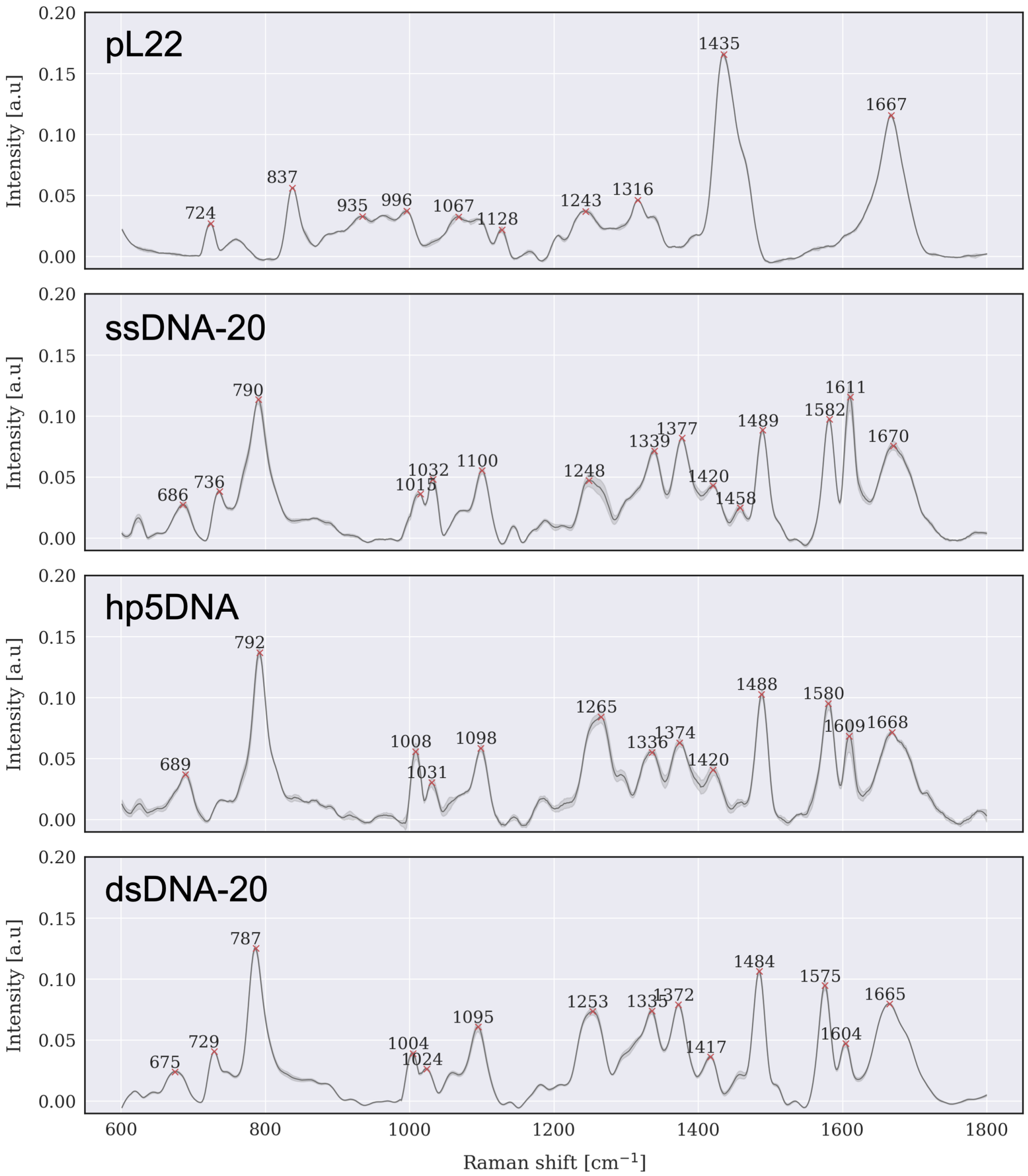


Figure S 8: *Average Raman spectra and deviations for dry samples of pL22 (n =3), ssDNA-20 (n =8), hp5DNA (n =6) and dsDNA-20 (n =6). Raw data are are processed and normalized as described in the main text (see Materials and Methods), with peak positions indicated in the plots.*

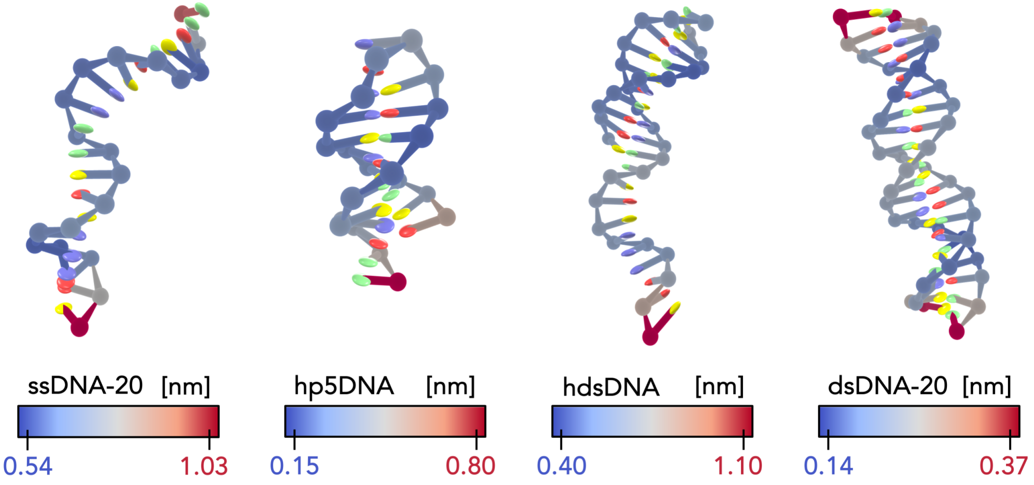


Figure S 9: *oxDNA centroid structures (from letf to right: ssDNA-20, hp5DNA, hdsDNA and dsDNA-20) with a Root Mean Square Fluctuation (RMSF) overlay over the backbone (blue-red color mapping). The RMSF is computed as the time averaged per-residue RMSD over the course of whole production simulation (see Materials and Methods).*

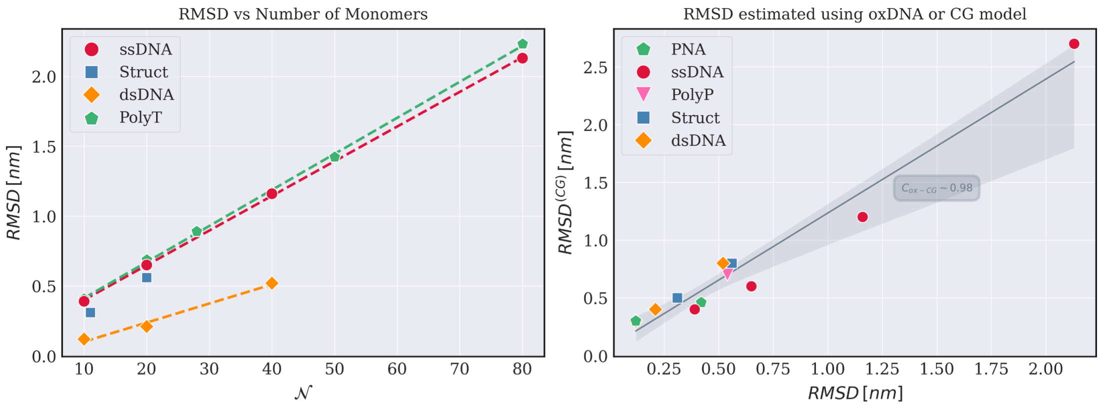


Figure S 10: *Left panel: Root Mean Square Deviation (RMSD) dependence on the number of monomers of several DNA strands, as estimated from oxDNA simulations; along with the ss and ds sequences, hp5DNA and hdsDNA are shown, together with a series of Poly-Thymine (PolyT) of different lengths. The dotted line is a fit to a line, showing how the scaling of the RMSD is clearly linear both in ss, ds and homopolymeric sequences. Right panel: correlation plot between the RMSD values estimated from oxDNA and from the CG model employed in this study; a linear regression with the relative interval region is shown, together with the Pearson correlation coefficient.*

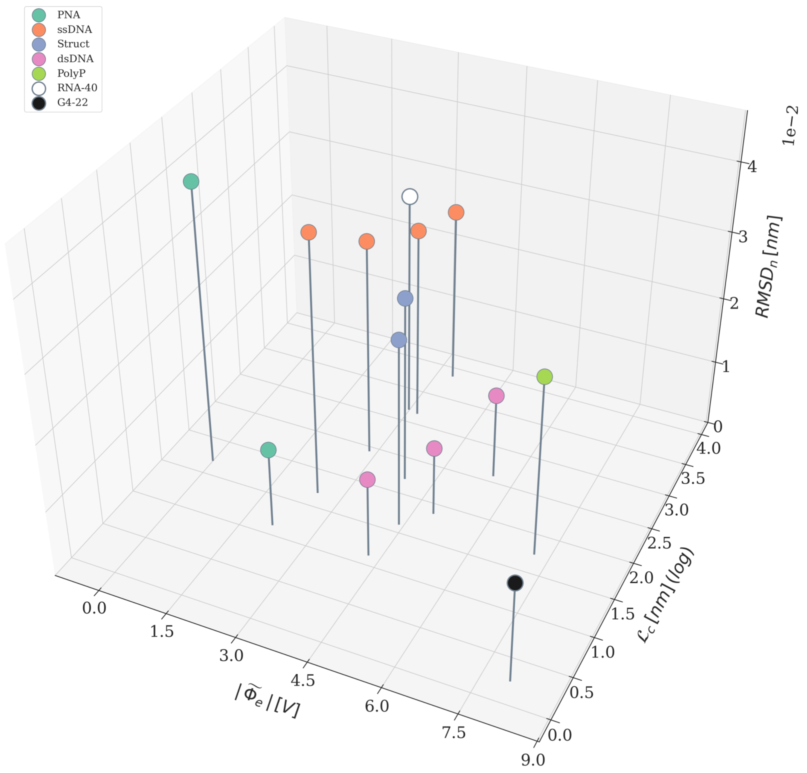


Figure S 11: *Comparison of the ONTs employed in this study with a G-quadruplex of 22 nucleotides (G4-22) and a RNA sequence of 40 nucleotides (RNA-40) in the parameter space of contour length $\mathcal{L}_c$, normalized average surface electrostatic potential $|\widetilde{\Phi}_e|$ and normalized $RMSD_n$. The color-coding refers to the different classes of sequences, with G4-22 and RNA-40 shown in black and gray, respectively.*

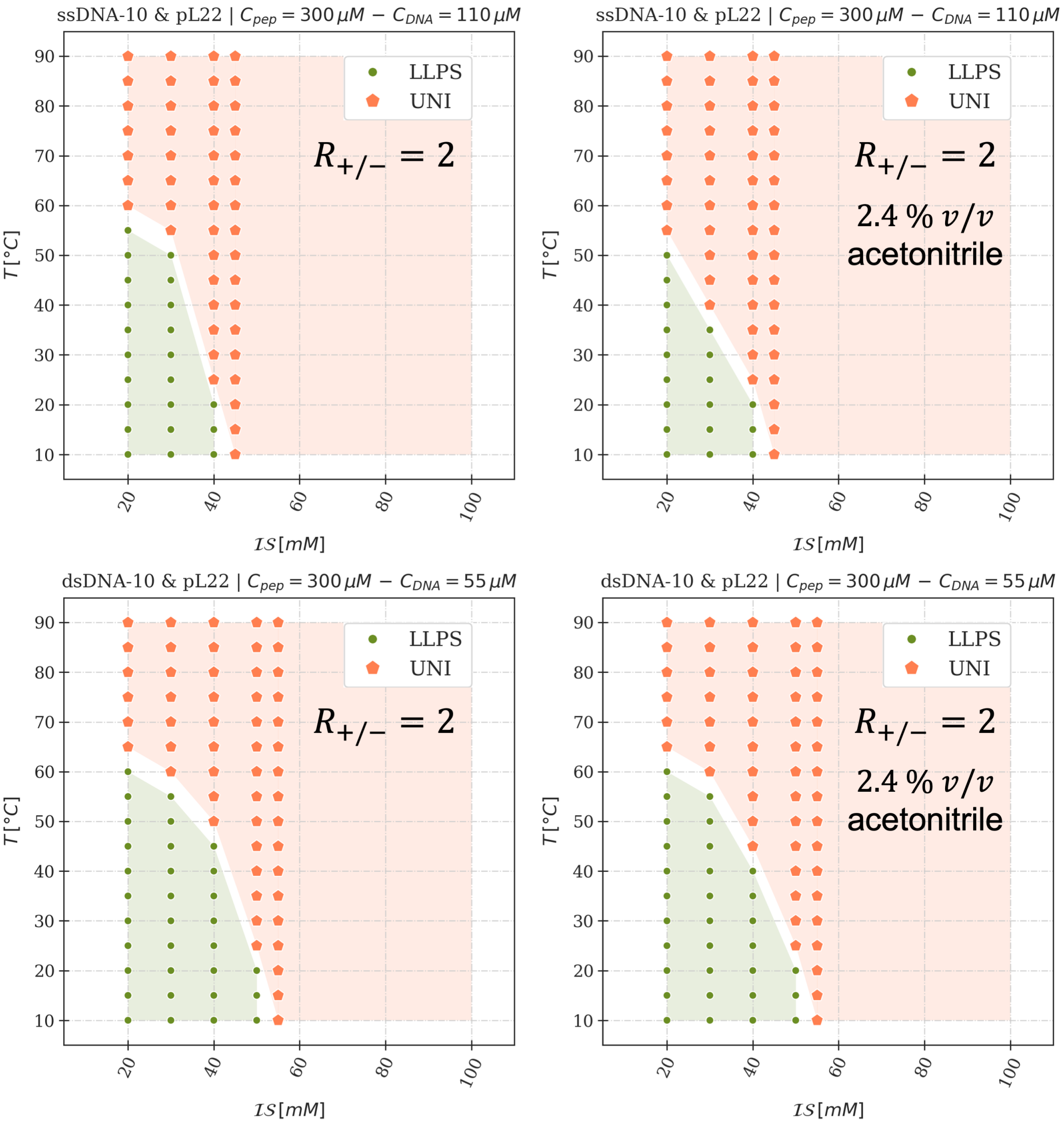


Figure S 12: *Experimental phase diagrams of mixtures of pL22 with ssDNA-10 (top row) and dsDNA-10 (bottom row). All phase diagrams refer to mixtures with charge ratio equal to 2 in water solution, 20 mM Tris-HCl pH 7.7 and added NaCl. Samples on the right column contained an additional 2.4% v/v acetonitrile, which was added to obtain the same solvent conditions used for PNA-10 hybrid. Monophasic and biphasic regions are denoted by the color and shape of markers and shaded regions (orange pentagons and green circles, respectively).*

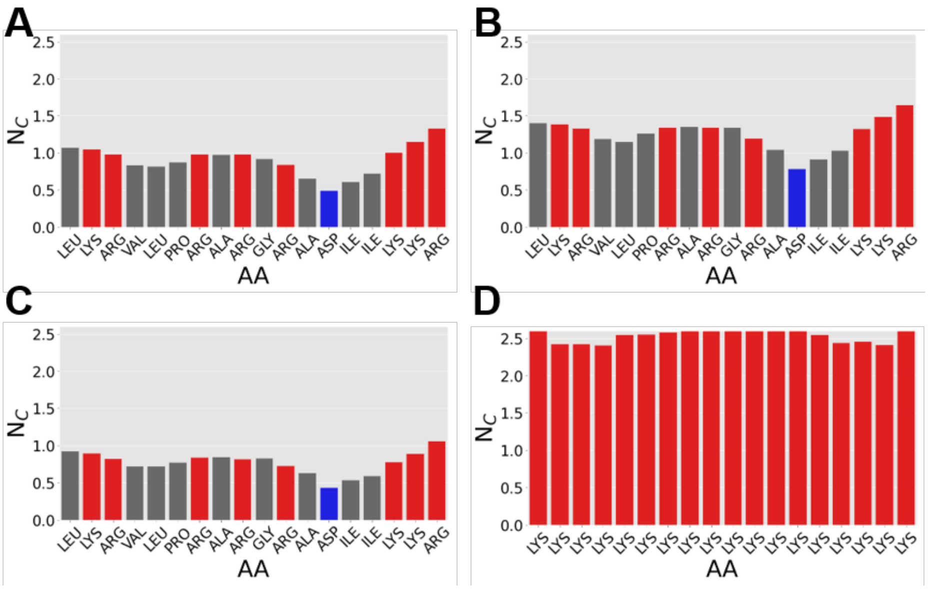


Figure S 13: *Average number of contacts with nucleotides, $N_c$, in simulations at $T$=303 K for the peptide residues inside the coacervates. A) pL22/ssDNA-20, $\mathcal{IS}$=75 mM (isotropic); B) pL22/dsDNA-20, $\mathcal{IS}$=75 mM (LC); C) pL22/dsDNA-20, $\mathcal{IS}$=225 mM (isotropic); D) polyK-18/dsDNA $\mathcal{IS}$=75 mM (solid).*

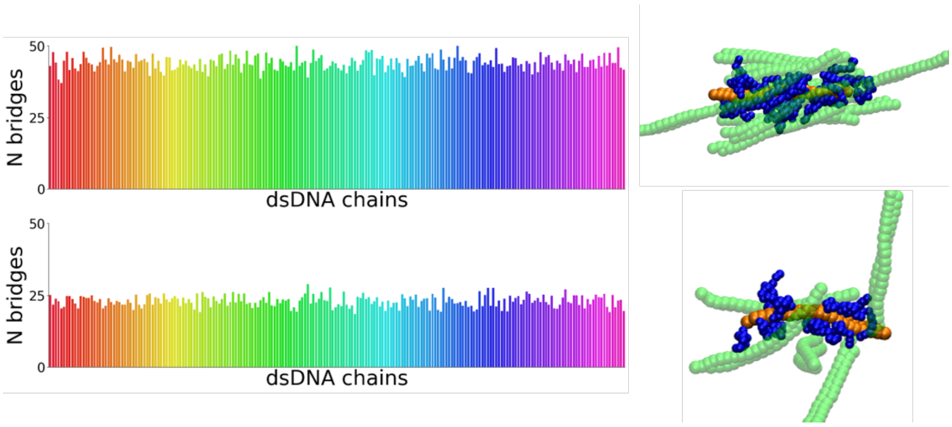


Figure S 14: *Average number of inter-DNA contacts bridged by peptide chains within pL22/dsDNA-20 coacervates (barplots), calculated for the individual DNA chains in the sample (from red to violet), and simulation snapshots showing one DNA reference chain (orange) surrounded by other DNA chains (green) bridged to it by peptides (blue). Simulations performed in the LC phase at $T$=303 K and $\mathcal{IS}$=75 mM (top row), and in the isotropic phase at $T$=303 K and $\mathcal{IS}$=225 mM (bottom row).*

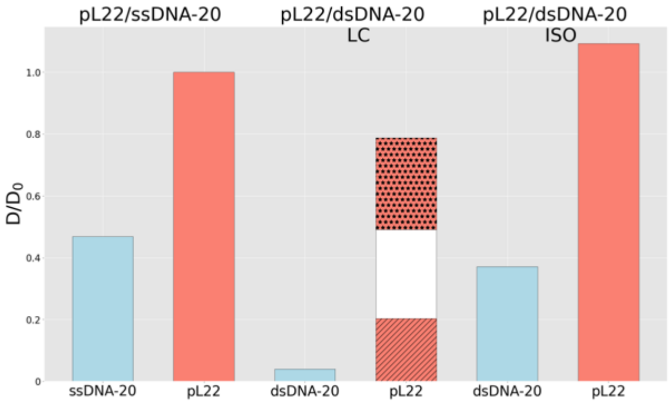


Figure S 15: *Translational diffusion coefficient of the center of mass of peptides (orange) and nucleotides (light blue), calculated from MD simulations at $T$ =303 K, for the dense phase of pL22/ssDNA-20 ($\mathcal{IS}$=75 mM) and of pL22/dsDNA-20 (LC at $\mathcal{IS}$=75 and isotropic at $\mathcal{IS}$=225 mM). In the bar for peptides in LC coacervates hatches are used to highlight the perpendicular (lines) and parallel (dots) contribution with respect to the director, while the white color is used to highlight the average diffusion coefficient value.*

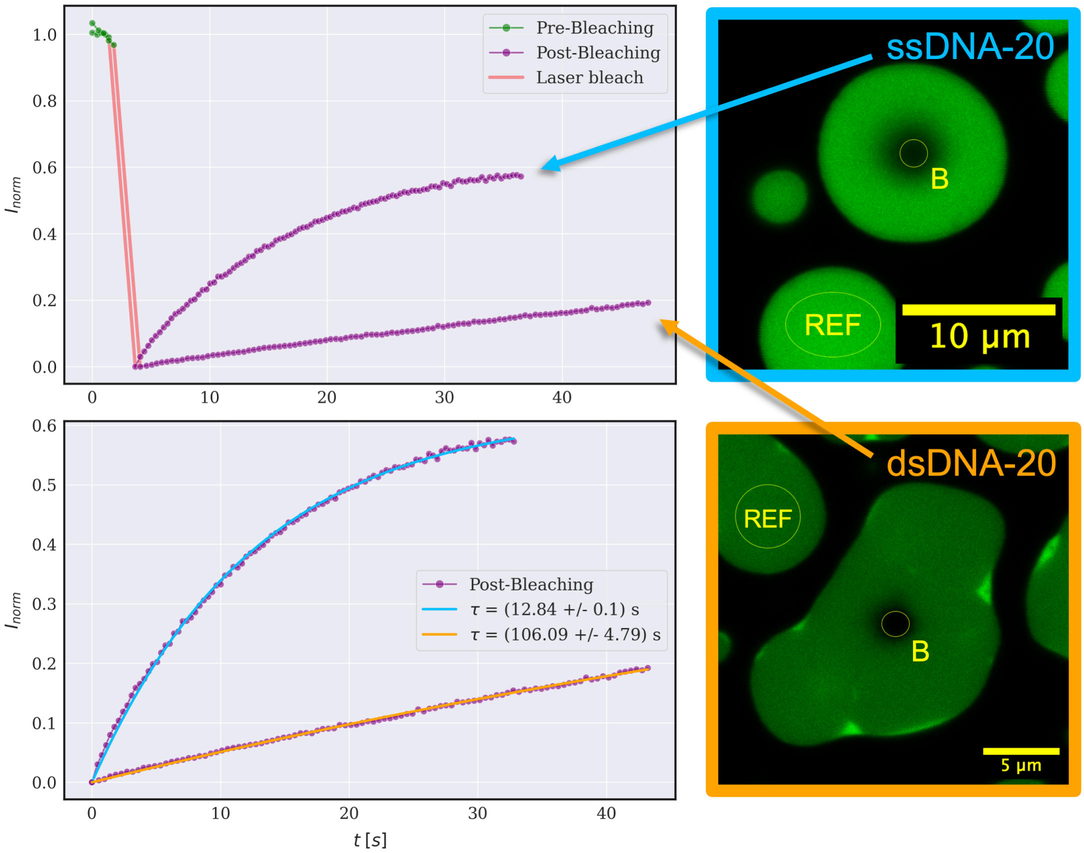


Figure S 16: *Fluorescence Recovery After Photobleaching (FRAP) experiment on mixtures of pL22 ($C_{pL22}$=300 µM) and ssDNA-20 (light blue micrograph, $C_{DNA}$=110 µM) or dsDNA-20 (orange micrograph, $C_{DNA}$=55 µM), with 1:200 FAM-labeled DNA strands. The two micrographs show the bleaching spots (yellow circular ROIs, B) and the regions used for photobleaching correction (yellow round ROIs, REF). The recovery curves on top show the normalized fluorescence signals for both samples prior and after laser irradiation, where intensities are processed as described in Material and Methods. The FRAP curve is reproduced on the panel below with the exponential recovery fit for both samples: the estimated characteristic times are $\tau \sim 13$ s for ssDNA-20 and $\tau \sim 106$ s for dsDNA-20.*

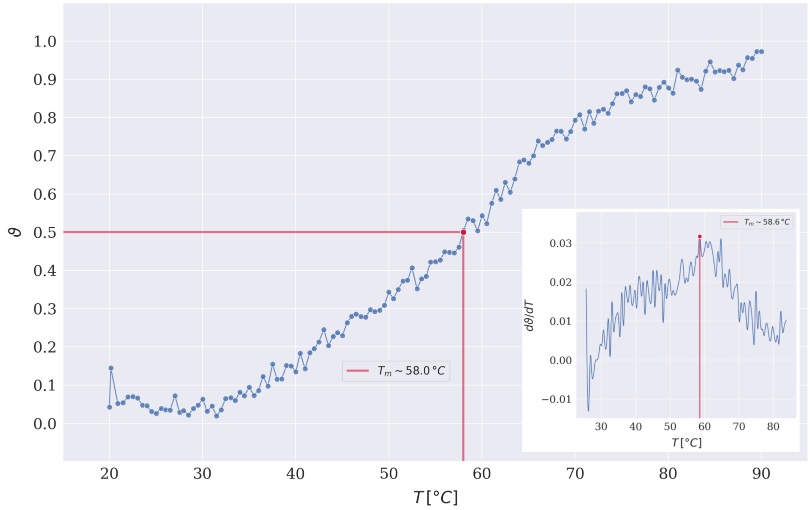


Figure S 17: *Normalized absorbance melting curve of PNA-10 hybrid at* $5\,\mu M$ *concentration in water solution, with a small* ($< 1\%$ $v/v$) *amount of acetonitrile. The absorbance spectra in the range 210-320 nm are collected and the raw signals around the 260 nm peak are then averaged and normalized: to a first approximation, the value* $\vartheta$=*0.5 provides the melting temperature (indicated by the red marker and lines), which is found to be around 58°C. The inset of the curve shows the numerical differentiation, upon smoothing and interpolation, with the maximum providing a more reliable estimate of the melting temperature.*

[11] Richard J Gowers, Max Linke, Jonathan Barnoud, Tyler John Edward Reddy, Manuel N. Melo, Sean L. Seyler, Jan Domanski, David L. Dotson, Sébastien Buchoux, Ian M. Kenney, , and O. Beckstein. MDAnalysis: a Python package for the rapid analysis of molecular dynamics simulations. In S. Benthall and

S. Rostrup, editors, Proceedings of the 15th Python in Science Conference, pages 98–105, Austin, TX, 2019.

[12] G. R. Luckhurst and G. W. Gray. The Molecular Physics of Liquid Crystals. Academic Press, New York, 1979.
